# Molecular basis of high-torque transmission of the *Vibrio* polar flagellar motor

**DOI:** 10.64898/2025.12.04.692373

**Authors:** Ling Zhang, Jiaxing Tan, Xuemin Duan, Xiaofei Wang, Michio Homma, Seiji Kojima, Yan Zhou, Yongqun Zhu

**Affiliations:** Department of Infectious Diseases of the Second Affiliated Hospital, School of Medicine, and School of Public Health, Life Sciences Institute, Zhejiang University, Hangzhou, China; The MOE Key Laboratory of Biosystems Homeostasis & Protection, and Zhejiang Provincial Key Laboratory of Cancer Molecular Cell Biology, Life Sciences Institute, Zhejiang University, Hangzhou, China; State Key Laboratory for Vegetation Structure, Function and Construction, Institute of Microbiology, College of Life Sciences, Zhejiang University, Hangzhou, China; Department of Physics, Nagoya University, Nagoya, Japan; Department of Biological Science, Graduate School of Science, Nagoya University, Nagoya, Japan; Shanghai Institute for Advanced Study, Zhejiang University, Shanghai, China; Cancer Center, Zhejiang University, Hangzhou, China

**Keywords:** polar flagella, flagellar motor, HT ring, hook, Leucine-rich repeat, *Vibrio*

## Abstract

The bacterial flagellar motors are huge protein nanomachines that drive rotation of the flagellum for bacterial motility, and have considerable diversity in structure among bacterial species, which enables the transmission of different torques to the flagellar filaments to propel bacteria and renders various swimming abilities. Relative to the bacterial peritrichous flagellar motors, the polar flagellar motors are the faster rotational machines and transmit high torque to drive bacterial high-speed motility in liquid and empower swimming in viscous environments. However, the structural basis of high-torque transmission of the polar flagellar motors is still unclear. Here we present an atomic-resolution cryo-electron microscope structure of the polar flagellar motor in complex with the hook from *Vibrio alginolyticus*, comprising 295 subunits from 18 proteins. Extensive inter-subunit interactions and additional phospholipids generate the higher rigidity of the rod. The LP ring utilizes more electrostatic charges on the inner surface and less physical contacts to facility the higher-speed rotation of the rod. The additional HT ring tightly binds to the outer surface of the LP ring to enhance the LP-ring stability. A previously function-unknown component FlrP enhances the interactions of 15 FliF peptides of the MS ring with the rod and stabilizes the LPHT and MS rings. The hook has two different states, L- and R-state, which provides structural flexibility of the hook to drive the unique forward-reverse-flick motility of the bacteria in the flicking process. This study provides unprecedented molecular insights into evolution and structural adaptions of the bacterial polar flagellar motors for high-torque transmission.

## Introduction

The majority of bacteria on Earth are polarly flagellated, possessing one or more flagella at the cell pole (*1–3*), such as in *Vibrio alginolyticus*, *Pseudomonas aeruginosa*, *Campylobacter jejuni*, and *Helicobacter pylori* (*1, 4–7*). The polar flagellar motor drives the rotation of the polar flagella for motility, which plays the key roles in bacterial environmental survival and pathogenicity (*8–10*). In contrast to the peritrichous flagellar motors that transmits low torque, such as ∼1,300 pN nm in *Escherichia coli* and *Salmonella* Typhimurium (*11–13*), the polar flagellar motors transmit high torque, like ∼3800 pN nm in *V*. *alginolyticus*, which renders the rotation of the polar flagellum at a remarkable speed of up to 1,700 revolutions per second (*11*) and enables bacterial swimming in viscous environments. We previously solved the cryo-electron microscope (cryo-EM) structure of the peritrichous flagellar motor of *S*. Typhimurium, and revealed its assembly mechanism by the C26-symmtric LP ring in the outer membrane, the MS ring in the inner membrane, the rod, the export apparatus, the cytoplasmic C ring, and the stator units that generate torque by transducing protons or sodium ions from the extracellular side to the cytoplasm (*14–20*). For high-torque transmission, the polar flagellar motor has evolved additional components and assembled a structure more complex than that of the peritrichous flagellar motor. For example, in *Vibrio* species, it contains additional H and T rings that surround the LP ring to form a LPHT ring in the outer membrane localization (*2, 3, 21–25*). Despite many progresses of cryo-electron tomography imaging and other structural efforts, the detailed structural adaption and molecular basis of high-torque transmission of the bacterial polar flagellar motors are still unclear. In addition, in *V*. *alginolyticus*, the polar flagellum drives the unique “forward-reverse-flick” motility pattern of the bacteria for cell reorientation, which differs significantly from the typical “run-and-tumble” pattern of the peritrichously flagellated bacteria (*1, 15, 26*). The flicking behavior of the bacteria requires a buckling instability process of the hook (*3, 27, 28*). The polar hook is usually straight to transmission the torque from the motor to the filament to push the bacterial cell body forward when the motor rotates in counterclockwise (CCW) or pull it backward once the rotational direction of the motor switches to clockwise (CW)(*1, 3*). However, in the flicking behavior, the straight hook bends under the thrust forces from the flagellar filament upon the change of the flagellar rotational direction from CW to CCW(*28, 29*) (*24, 28, 30, 31*). During the bending of the hook, the subunits of its component FlgE at the innermost side produce potential structural collapses. It is unknown how the straight hook obtains the structural flexibility to allow the bending and overcomes the potential collapses. In this study, we determined a cryo-EM structure of the polar flagellar motor-hook complex from *V*. *alginolyticus*, which provides unprecedent insights into the structural evolution for the high-torque transmission of the polar flagellar motor.

## Results

### Overall structure of the polar flagellar motor-hook complex

FlgL is an essential protein responsible for connecting the hook to the filament of the polar flagellum in *V. alginolyticus* (*2, 32*). To avoid the effects of the filaments and the lateral flagella during purification of the polar flagellar motor particles, we generated a Δ*flgL* mutant strain of *V. alginolyticus* KK148 that could not produce the lateral flagella but was able to assemble multiple polar flagella at the cell pole(*32–34*). The particles of the polar flagellar motor-hook complex of *V. alginolyticus* were successfully purified from the Δ*flgL* mutant strain (*14*) and contained the LPHT ring, the MS ring, the rod, the export apparatus and the hook (Fig. 1a and Fig. S1a). We also disassembled the purified particles of the motor-hook complex by reducing the concentration of the salts in the purification buffer to obtain the particles of the LPHT ring, which are highly stable even after the motor disassembly, for verifying its structural symmetry individually during structure determination (Fig. S1a, b).

**Figure 1.**
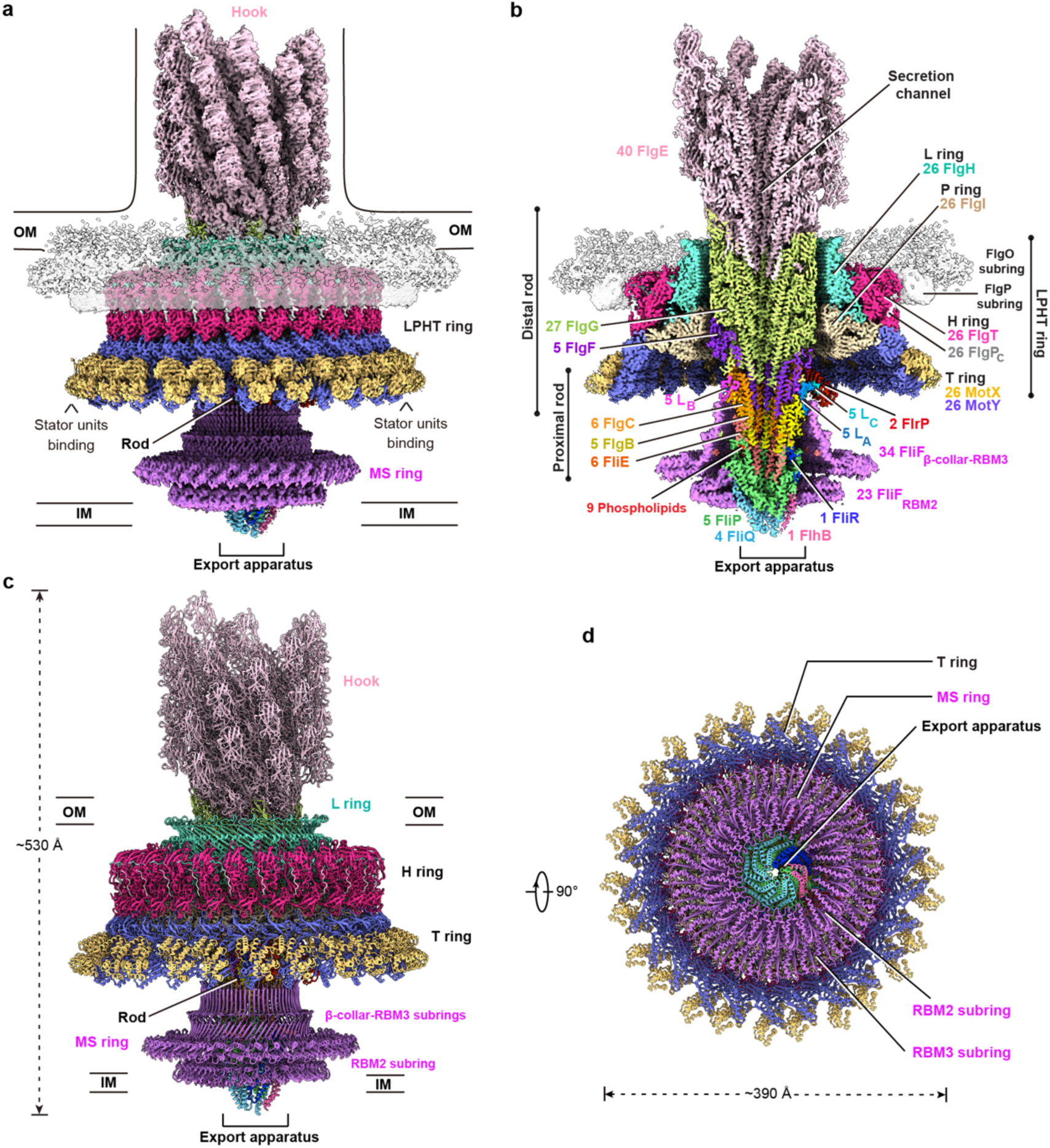
Overall structure of the polar flagellar motor-hook complex of *V. alginolyticus*. (a and b) The side (a) and cross-section (b) views of the merged density map of the polar flagellar motor-hook complex after local refinements. The map was obtained by fitting the high-resolution density maps of six locally refined regions, including the LPHT ring, the hook (L-type), the distal rod, the proximal rod with the export apparatus, the β-collar-RBM3 subrings (C34 symmetry) and the RBM2 subring (C1 symmetry) of the MS ring into the global density map of the polar flagellar motor-hook complex reconstructed with C100 symmetry. Semi-transparent illustration of the densities of the FlgP and FlgO subrings are applied to show the overall structure of the LPHT ring. The subunit numbers of the components are labeled as indicated (b). (c and d) The side (c) and bottom views (d) of the final structural model of the polar flagellar motor-hook complex. OM, outer membrane; PG, peptidoglycan; IM, inner membrane.

Due to its large box size of the overall three-dimensional (3D) volume of the whole motor-hook complex, we carried out reconstruction with C100 symmetry and successfully obtained a clear density map of the whole complex, which allowed us to assemble our locally refined density maps of different parts of the complex into the volume for overall structural modelling (Fig. S1c). Two-dimensional (2D) classification of the end-on viewed LPHT ring particles revealed a 26-fold symmetry of the LPHT ring (Fig. S1d). Further 3D classification of the particles of the LPHT ring revealed an unexpected 13-fold symmetry in its bottom region, indicating a mismatch between the 26- and 13-fold symmetries within the LPHT ring (Fig. S1c). Reconstruction with C13 symmetry yielded a density map of the LPHT ring with a resolution of ∼2.95 Å, which allowed the *de novo* model building of each subunit (Fig. S1c, S2). There are two layers of ring-like and smeared densities at the top region of the H ring, which could not be structurally modelled and are possibly the FlgO subring and the lipid-bound region of the FlgP subring, respectively, according to the previous cryo-electron tomography (cryo-ET) imaging analyses(*21, 22*) (Fig. 1a, b).

2D classification analyses revealed the hook in the complex particles was in two different conformational states (Fig. S1c), which are hereafter names as the L- and Rtype, respectively, and characterized with the left- and right-handed helical arrangements of the D2 domains of the FlgE subunits at the outermost layer. The density maps of the L- and R-hooks were reconstructed to ∼3.2 Å and ∼3.4 Å, respectively (Fig. S1c, S2). Local refinement of the MS ring without imposed symmetry produced a ∼3.6 Å density map and revealed the 34-fold and 23-fold symmetries in the upper β-collar-RBM3 and lower RBM2 subrings, respectively (Fig. S1c, S2). The density map of the β-collar RBM3 subrings of the MS ring was finally reconstituted to ∼2.9 Å with C34 symmetry (Fig. S1c, S2). Focused refinement also produced high-resolution density maps for other regions in the complex, including the axis region containing the rod, the export apparatus and a part of the hook, the proximal rod region with the export apparatus, and a middle part of the rod, to the resolutions of ∼3.0 Å, ∼3.0 Å and ∼3.5 Å, respectively (Fig. S1c, S2).

The final model of the polar flagellar motor-hook complex contains 295 subunits from 18 proteins, including 26 subunits of each component of FlgH, FlgI, FlgT, MotX, MotY and FlgP in the LPHT ring; 34 FliF subunits in the MS ring; 1 FlhB, 4 FliQ, 5 FliP and 1 FliR subunits in the export apparatus; 6 FliE, 5 FlgB and 6 FlgC subunits in the proximal rod; 5 FlgF and 27 FlgG subunits in the distal rod; 40 FlgE subunit in the part of the hook; and 5 previously function-unknown leucine-rich repeat (LRR) proteins (hereafter named as FlrP) that are bound onto the middle region of the rod (Fig. 1a-c). In the complex, the LPHT ring surround the distal rod, while the MS ring wraps the export apparatus and the proximal rod (Fig. 1a, b). The whole structural model of the polar flagellar motor-hook complex has a molecular weight of ∼10 MDa with an outermost diameter of ∼390 Å at the T ring and a height of ∼530 Å. The molecular weight and size of the polar flagellar motor we obtained significantly exceed those of the *Salmonella* peritrichous flagellar motor (Fig. 1c, d).

### The additional HT ring enhances the stability of the LP ring

In the motor-hook complex, the LP ring consists of 26 FlgH and 26 FlgI subunits with an inner diameter of ∼130 Å to embrace the distal rod (Fig. 2a). Each FlgH subunit interacts with the prior and next three FlgH protomers, while each FlgI subunit interacts with the prior and next two FlgI protomers (Fig. S3). FlgH harbors a distinctive N-terminal loop (L_N_ loop) that has a sequence much longer than that in the peritrichous FlgH and adopts a more compact conformation in its middle and C-terminal regions (Fig. 2a, b, and Fig. S3a-d), leading to a 1-versus-3 relatively looser interaction manner between the FlgH and FlgI subunits at the L-P interface, rather than the 1-versus-4 manner in the peritrichous flagellar motor (Fig. S3g). The N-terminus of the L_N_ loop extends outward to interact with the FlgT and FlgP subrings of the H ring (Fig. 2b, e and Fig. S5c).

**Figure 2.**
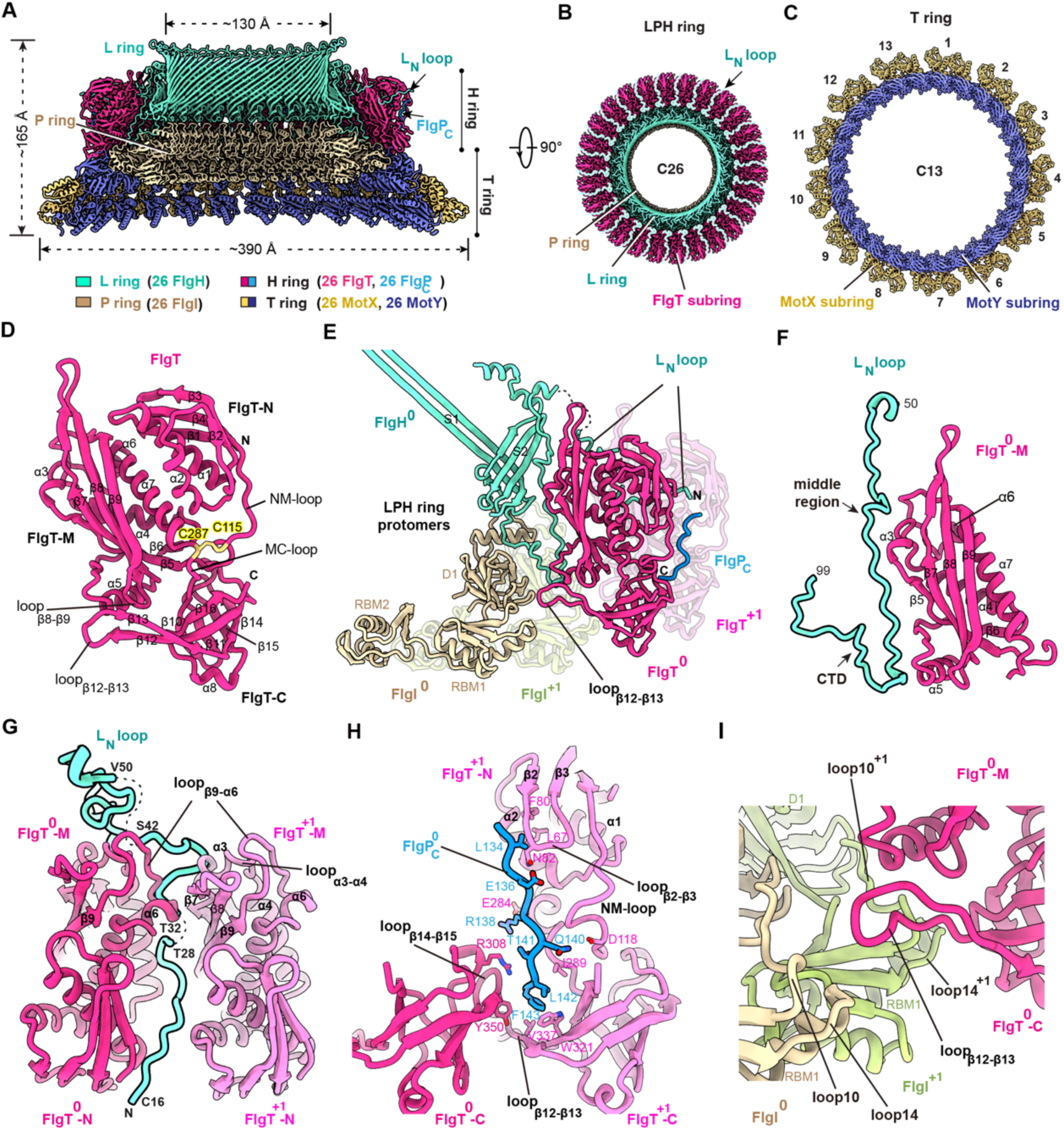
The structure of the LPHT ring and its assembly. (a) Cross-section view of the structure of the LPHT ring in the polar motor-hook complex. (b) Top view of the structure and C26 symmetry of the LPH ring. (c) Top view of the structure and C13 symmetry of the T ring. (d) Structure of FlgT in the LPHT ring. (e) Interactions of FlgT with the subunits of FlgH, FlgI and FlgP_C_ in the LPHT ring. (f and g) Interactions of the middle (f) and N-terminal regions (g) of the L_N_ loop of FlgH with the FlgT subunits in the LPHT ring. The flexible regions (residues 29-31 and residues 43-48) of the L_N_ loop are indicated with dashed lines. (h) Detailed interactions of FlgP_C_ with the FlgT subunits. (i) Interactions between the FlgT subring and the P ring.

The H and T rings surround the LP ring and consist of 26 FlgT and 26 FlgP, and 26 MotX and 26 MotY subunits, respectively (Fig. 2a-c). In the H ring, FlgT consists of three domains, FlgT-N, FlgT-M and FlgT-C(*35*), and adopts a closed conformation, which is different from the open conformation in apo crystal structure(*35*) (Fig. 2d and Fig. S5a, b). Different to the LP and H rings, which have C26 symmetry, the T ring adopts the C13 symmetry(*22*) (Fig. 2b, c). Each protomer of the T ring contains two MotX-Y heterodimers, MotX_1_-Y_1_ and MotX_2_-Y_2_ (Fig. S4b-e and S5d), due to the two different conformations of the C-terminal regions of MotX, which is essential for the interactions of the two MotX-Y dimers in the protomer. The C-terminal region of MotX_2_ extends out from its main body and inserts into a hydrophobic cavity in MotX_1_-C (Fig. S4f and Fig. 3g). In contrast, the C-terminal region of MotX_1_ folds back into a cleft between α4 of MotY_1_-C and α9 of MotX_1_ in the MotX_1_-Y_1_ dimer (Fig. S4e). In the T ring, the N-terminal domains of MotY (MotY-N) interact with each other in a head-to-tail manner (Fig. S4b, h). MotX also interacts with the upper two MotY-N domains to bridge the two subrings of the T ring (Fig. S4j).

**Figure 3.**
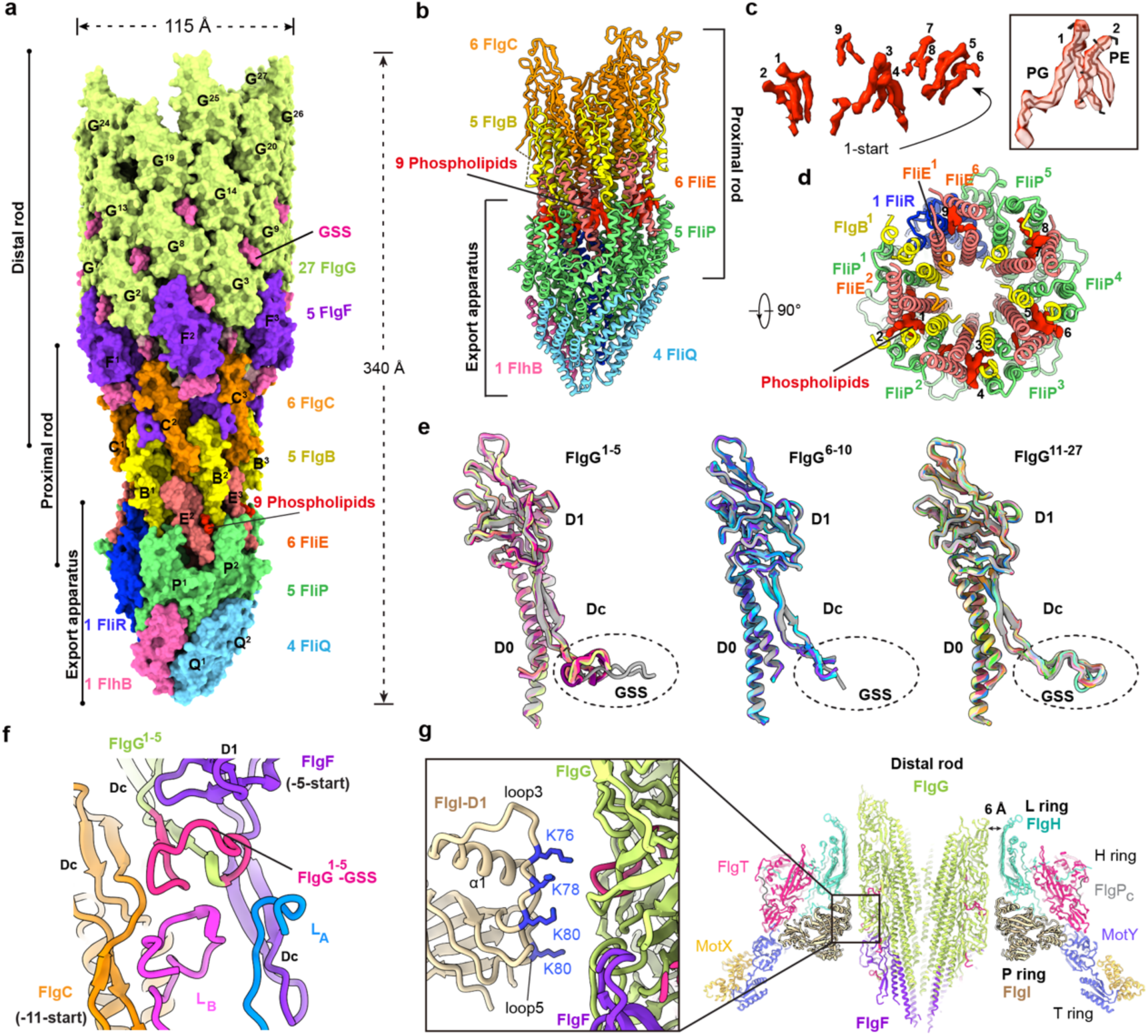
The structure of the rod. (a) Overall structure of the rod with the export apparatus in the polar flagellar motor-hook complex. The rod and the export apparatus are represented in surface. (b and c) Side (b) and top views (c) of the binding of nine phospholipids into the grooves at the interface of the proximal rod and the export apparatus. (d) Cryo-EM density maps of nine phospholipid molecules. The densities of the nine phospholipids are colored in red (b, c, and d). Close-up view of the first and second phospholipids are shown at right and modeled as PG and PE molecules, respectively. (e) Structural comparison of the FlgG subunits in the rod. (f) Interactions of the GSS regions of the FlgG^1-5^ subunits with the adjacent subunits in the rod. (g) Detailed interactions between the LPHT ring and the rod. Lysine residues of the D1 domain of FlgI, which are involved in the interactions with the rod, are shown as sticks and colored in blue.

The HT and LP rings generate extensive interactions. The L_N_ loops of FlgH clamp the FlgT-M domains in the H ring (Fig. 2a, b, e, g). The middle region of L_N_ binds to α3, α5, the C-terminus of β5, loop_α3-α4_, loop_β6-α5_, and loop_β9-α6_ of FlgT-M (Fig. 2f). The T32-S52 fragment at the N-terminal region of L_N_ extends outwards and inserts into a groove formed by α6 and loop_β9-α6_ of FlgT-M with the loop_α3-α4_ and β7 of FlgT^+1^-M (Fig. 2g). The proximal region (C16-T28) of L_N_ binds into the deep cleft between the adjacent FlgT-NM domains (Fig. 2g). The invisible lipid-modified N-terminus of the L_N_ loop likely functions as an anchor with the FlgP subring to anchoring the LPHT ring to the outer membrane (Fig. S5c). The C-terminal tail (Q133-F143) of each FlgP subunit obliquely binds into an open groove in the FlgT subring (Fig. 2h). In addition, the loop_β12-β13_ of FlgT-C interact with the loops 10 and 14 of FlgI at the interface of the H ring with the LP ring (Fig. 2i). MotY-N also inserts into the cleft between the FlgI-RBM1 and FlgT-C domains to stabilize the P ring (Fig. S4i).

### Phospholipids and more inter-subunit interactions increase the rod rigidity

Different from the 24 FlgG subunits in the *Salmonella* peritrichous rod, there are 27 subunits of FlgG of *V. alginolyticus* in the distal rod of the polar rod (Fig. 3a). The polar rod is located on the export apparatus, which is bound by two N-terminal transmembrane helices of FlhB from the bottom (Fig. 3a, b and Fig. S6a). The export apparatus adopts an expanded configuration to accommodate the subunits of the proximal rod (Fig. 3b). There are nine phospholipid molecules, which are possibly phosphatidylglycerols and phosphatidylethanolamines, at the interface of the export apparatus and the proximal rod (Fig. 3b-d). These phospholipids insert into the clefts formed by FliE^2^-FlgB^2^-FliP^2^, FliE^3^-FlgB^3^-FliP^3^, FliE^4^-FlgB^4^-FliP^4^, FliE^5^-FlgB^5^-FliP^5^ and FliR-FliE^6^-FliE^1^-FlgC^1^, respectively (Fig. 3b, d) and act as glue to strengthen the interactions between the export apparatus and the proximal rod, thereby highly increasing the rigidity of the proximal rod.

The subunits of the polar rod form more inter-subunit interactions than those in the peritrichous rod. Each polar rod protein harbors unique structural features (Fig. S6d-g). FliE contains an elongated α2 helix for interaction with the export apparatus (Fig. S6d). The N-termini of the α_N_ helices of the FlgB subunits twist to interact with the upper regions of the FliP subunits, thereby enhancing interactions between the export apparatus and the proximal rod (Fig. S6e). FlgC has a unique lip structure to interact with the FliF peptide loops and the newly identified LRR proteins FlrPs (details are discussed below) (Fig. S6f). In contrast to the *Salmonella* FlgF, FlgF of the polar rod lack an α3 helix (Fig. S6g). The FliG subunits in the polar rod can be classified to three classes, FliG^1-5^, FliG^6-10^, and FliG^11-27^, based on the conformations of their FlgG-specific sequence (GSS) regions (Fig. 3e). The GSS regions of FlgG^1-5^ adopt a compressed conformation and extensively interact with the FlgC^1-5^- Dc and FlgF^1-5^-DcD1 domains, as well as with the FliF peptide loops, L_A_ and L_B_, along the -11- and -5 start directions (Fig. 3e, f). The GSS regions of FlgG^6-10^ are unfolded in the structure, while the FlgG^11-27^ GSS region interacts extensively with the neighboring FlgG and FlgF subunits in the -16, -10, and -5 directions to enhance the rigidity of the distal rod (Fig. S6h). FlgG^11^ also interacts with FlgC^6^ at the -5-start position (Fig. S6h).

### Stronger electrostatic interactions at the rod-LP ring interface

At the LPHT ring-rod interface, only the P ring physically contacts the D1 domains of FlgF^1-5^, the D1 domains of FlgG^1-10^ and the GSS regions of FlgG^14-18^ via the lysine residues, K76, K78, and K80 in loop3 and K112 in loop5, of the FlgI-D1 domains (Fig. 3g). In contrast to two lysine residues and five polar residues including glutamine and asparagine in FlgI-D1 of the P ring to form a hydrogen-bond ring with the rod in the peritrichous flagellar motor(*14*), four rod-interacting lysine residues in the FlgI-D1 domain in the polar flagellar motor result in fewer physical contacts but stronger electrostatic interactions between the rod and the LPHT ring (Fig. 3g). In addition, the distal rod surface carries a greater abundance of negatively charged residues with nine aspartates and ten glutamates on each D1 domain of the FlgG subunits, in contrast to six aspartates and five glutamates on that of the peritrichous flagellar motor (Fig. S6i-l). The L ring in the polar flagellar motor also carries more negatively charged residues with six aspartates and two glutamates on the rod-facing surface of each FlgH subunit, rather than two aspartates and one glutamate on each FlgH surface of the peritrichous flagellar motor (Fig. S6i-k), suggesting that the LP ring utilizes stronger electrostatic interactions and less physical contacts to stabilize the rod during the rotation at the higher speed.

### More rod-binding FliF loops from the MS ring

Like that in the peritrichous flagellar motor, the MS ring in the polar motor has the symmetry mismatch between the upper 34-fold symmetric RBM3-β-collar subring and the lower 23-fold symmetric RBM2 subring (*25*) (Fig. S7a, b). The MS ring has the outermost diameters of 230 Å at the RBM3 subrings (Fig. 4a). The lower RBM2 subring binds to the export apparatus via the two long loops, loop 1 and loop 2, of each RBM2 domain (Fig. S7c). The RBM3-β-collar subring encompasses both the upper region of the export apparatus and the lower region of the proximal rod (Fig. 4a). The N-terminal α1 helices of the six FliE subunits and the Dc loops of the five FlgB subunits, which are extended outward from the proximal rod, bind to the inner surface of the β-collar subring of the MS ring (Fig. 4a).

**Figure 4.**
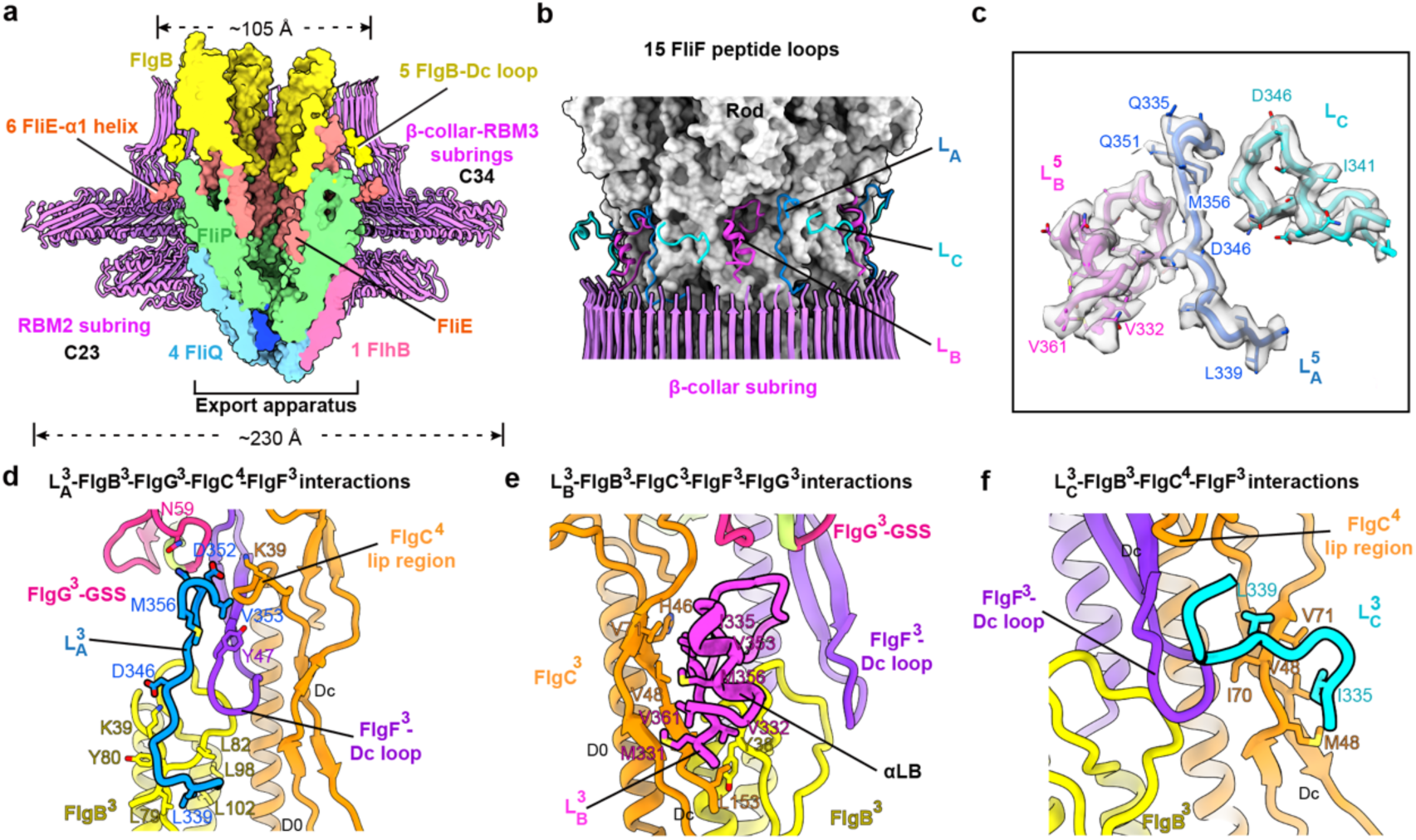
The MS ring-rod interactions and the FlrP protein. (a) Structure of the MS ring and its interactions with the proximal rod and the export apparatus. (b) Interaction of the 15 FliF peptide loops with the rod. The rod is represented in surface. the L_A_, L_B_ and L_C_ loops of FliF are colored in blue, purple and cyan, respectively. (c) Representative cryo-EM densities of L_A_, L_B_ and L_C_. The L_A_^5^ loop, L_B_^5^ loop and L_C_^5^ loop are illustrated. (d, e, and f) Detailed interactions of the L_A_ loop (d), L_B_ loop (e) and L_C_ loop (f) with the rod. The interaction of the L_A_^3^ loop, L_B_^3^ loop and L_C_^3^ loops with the rod are illustrated as indicated.

Locally refined maps of the proximal rod and the middle rod revealed that there are 15 FliF peptide loops, which extend upward from the β-collar subring of the MS ring, to bind the rod for torque transmission (Fig. 4b), in contrast to 11 extended FliF peptide loops in the peritrichous flagellar motor(*14*). The 15 FliF peptide loops are grouped into five sets, each of which comprises an L_A_, L_B_ and L_C_ loop and is packed helically on the surface of the rod (Fig. 4b, c and Fig. S7d). The L_A_ loop is bound into the long clefts formed by the upper region of FlgB^1-5^-Dc domain with the FlgF^1-5^-Dc, FlgC^2-6^-lip, and FlgG^1-5^-GSS regions at the 11-, 6- and 16-start positions (Fig. 4d). The L_B_ loop, which harbors a unique α_LB_ helix, inserts to an open pocket generated by the upper region of FlgB^1-5^-Dc, the D0 and D1 domains of FlgC^1-5^ and FlgF^1-5^, and the FlgG^1-5^-GSS at the -11-, 5-, 11- and 16-start positions, respectively (Fig. 4e). The L_C_ loop interacts with the concave surface on the lower region of FlgF^1-5^-Dc and the middle region of FlgC^1-5^-Dc (Fig. 4f). The N-terminus of L_C_ interacts with the α_LB_ helix of L_B_^+1^ of the next set, while the C-terminus of L_C_ contacts with the L_A_ loop and the lip region of FlgC^1-5^ (Fig. 4b), thereby reinforcing the inter-loop interactions of the 15 FliF peptide loops on the surface of the rod.

### A previously unknown component in the polar flagellar motor

In addition to the 15 FliF loops, a 3.5 Å-resolution local density map of the proximal rod revealed that there are five β-solenoid-like structures surrounding the rod (Fig. S1c). Mass spectrometry analysis revealed that there was a previously unidentified protein of 21.5 kDa (protein ID: BCB49735.1, named as FlrP) in the purified particles of the polar flagellar motor-hook complex (Fig. S8a). FlrP contains a Sec signal peptide at its N-terminus (residues M1-A19) and is a typical leucine-rich repeat (LRR) protein. Western blotting using a monoclonal antibody specific to FlrP also confirmed its presence in the polar flagellar motor (Fig. S8b). Among the five FlrP subunits, FlrP^1^ and FlrP^5^ have very clear densities, which enabled us to build all residues in its structure (Fig. 5a, b and Fig. S8c). The structure of FlrP consists of a β-solenoid LRR repeat core with an N-terminal head (Y20-Y34) and a C-terminal tail (G195-P199) (Fig. 5b and Fig. S8d). Structural analysis of FlrP^1^ and FlrP^5^ revealed that the FlrP subunits not only bind to the rod, but also interact with the L_B_ and L_C_ loops of the FliF subunits to strengthen the MS ring-rod interactions (Fig. 5d-g). The head regions of FlrP^1^ and FlrP^5^ bind into the clefts between the lip and cap regions of FlgC^2^ and FlgC^6^, and interact with the GSS regions of FlgG^2^ and FlgG^6^ and the D1 domains of FlgF^2^ and FlgG^1^, respectively (Fig. 5d-g). FlrP^1^ and FlrP^5^ also interact with the L_C_^1^ and L_C_^5^ loops, respectively, through the coil 4-6. FlrP^1^ interacts with α_LB_ of the L_B_^2^ loop through the coil 1-2 (Fig. 5d-g).

**Figure 5.**
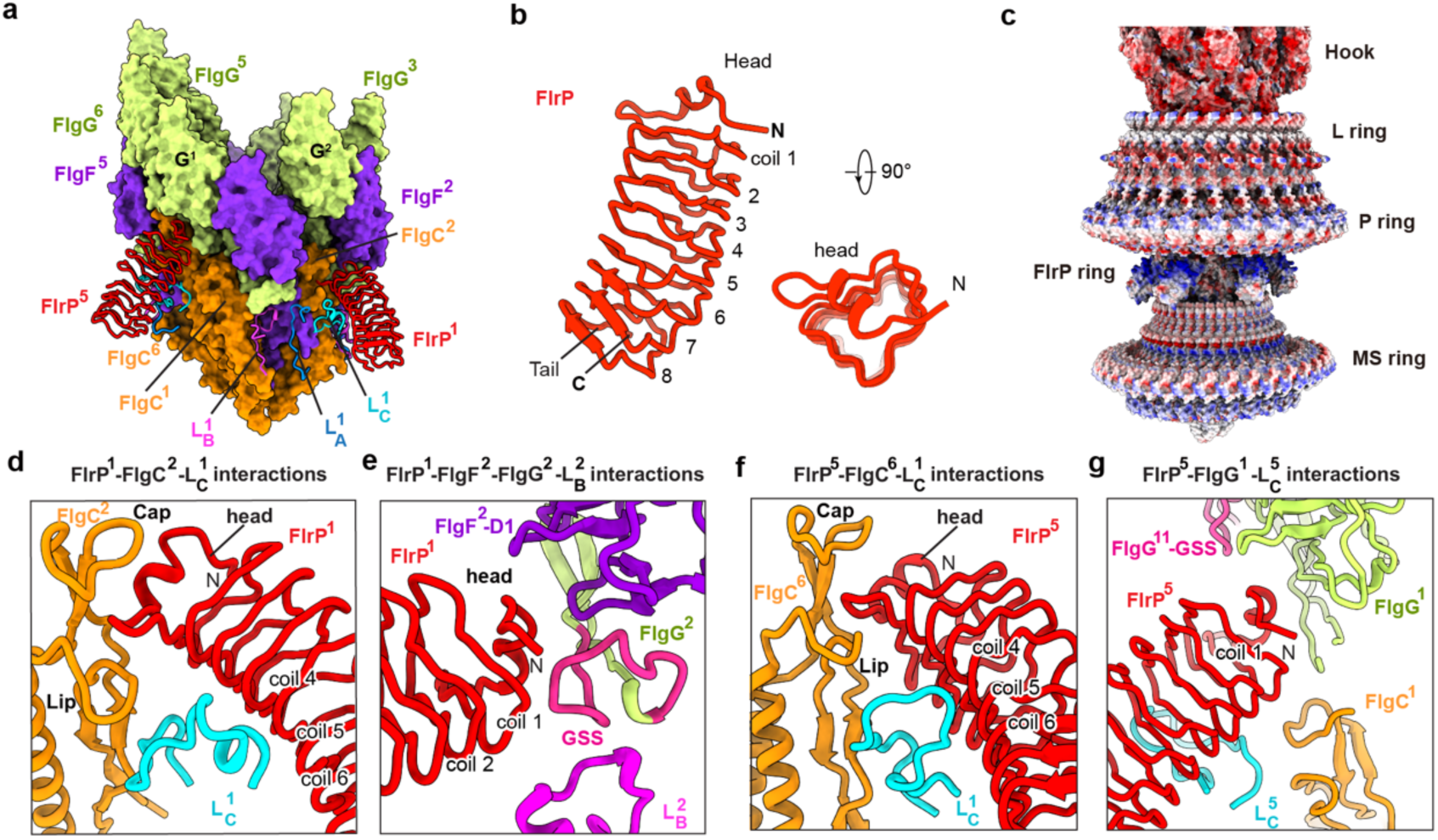
The FlrP protein. (a) Interactions between the FlrP^1^ and FlrP^5^ subunits with the rod. (b) Structure of FlrP. (c) Electrostatic potential distribution of the hook, the LP ring, the FlrP subunits and the MS ring in the polar flagellar motor. (d-g) Detailed interactions of the FlrP subunits with the rod and the rod-binding loops of FliF. The interactions of the FlrP^1^ (d and e) and FlrP^5^ (f and g) subunits with the rod and the L_B_ and L_C_ loops are illustrated as indicated.

On the surface of the rod, the five FlrP subunits are helically packed with the same helical parameters of the rod between the LPHT ring and the MS ring, and form a spiral ring with an outer diameter of ∼160 Å, which is much larger than the inner diameters of the RBM2 subring (∼145 Å) and the β-collar subring of the MS ring (∼105 Å) (Fig. S8e, f). The upper surface of each FlrP subunit is highly positively charged with the residues K54, K66, K69, R88, K92, K94, R109, K112, K130, R132, and K134, and produce electrostatic repulsion to the inner lower surface of the P ring that is also positively charged (Fig. 5c and Fig. S8g-i), suggesting that the FlrP subunits not only stabilize the FliF loop-rod interactions, but also likely act as charged spacers to maintain the positions of the LPHT and MS rings during the high-speed rotation of the polar motor (Fig. 5c). Consistent with the important role of the FlrP subunits in the structure, gene deletion of *flrP* significantly reduced the motility of *V. alginolyticus* (Fig. S9a). FlrP is also conserved in 13 species of the *Vibrionaceae* family, including *Vibrio*, *Photobacterium*, *Aliivibrio*, *Salinivibrio*, *Grimontia* and *Enterovibrio* (Fig. S9c). Its homologues have similar structures and harbor highly conserved the rod- and L_B_/L_C_- interacting residues (Fig. S9d). Gene deletion of the FlrP homologue in *V. parahaemolyticus* also reduced the motility of the bacteria (Fig. S9b). In contrast to most of the flagellar genes that are generally located on chromosome I (*36, 37*), the *flrP* gene is located on chromosome II of *V. alginolyticus* and *Vibrio* species, indicating a conserved new regulation mechanism between chromosomes in the polar flagellar system.

### Two types of the hook

In contrast to the curved hook in the peritrichous flagellum (*15*), cryo-EM imaging analysis revealed the presence of two types of the straight hook, L- and R-type, in the particles of the polar motor-hook complex (Fig. 6a, b and Fig. S1a, c). About 63.2% of the particles harbor the L-type hook, while about 36.8% have the R-type hook (Fig. S1c). In both types of the hook, the FlgE subunits of *V. alginolyticus* assemble helically along the 1-start helical direction in a right-handed manner, and generate a three-layered tubular structure through the D0, Dc-D1 and D2 domains, respectively (Fig. 6a, b). The structures of the inner and middle tubes of the L-type hook are the same as those in the R-type hook (Fig. 6a, b). However, relative to the L-type hook, the outer tube of the R-type hook is in compressed conformation (Fig. 6a, b and Fig. S10g). In the R-type hook, the D2 domains of the FlgE subunits (R-FlgE) pack via a head-to-tail interaction manner along the 6-start direction in the outer tube, which resembles the assembly of the D2 domains of the *Salmonella* FlgE subunits in the peritrichous curved hook(*14*) (Fig. 6a, c and Fig. S10a), whereas in the L-type hook, the D2 domains of the FlgE subunits (L-FlgE) are packed along the 11-start direction with the same head-to-tail manner (Fig. 6b, f), suggesting that the R-type hook likely functions as the hook of *S*. Typhimurium and represents the state of the hook under the thrust force generated by the filament during the CCW rotation of the polar flagellar motor and the forward swimming of the bacteria, while the L-type hook represents the state of the hook under the pulling force from the filament upon the rotation of the motor in the CW direction to pull the bacterial cell backward. Different to the *Salmonella* FlgE, FlgE of *V. alginolyticus* has evolved a longer conserved tip region (L46-T52) in the Dc domain (Fig. S10a) and harbors unique loop_β3-β4_ and loop_E_ in the D1 domain, both of which are in the lifted conformations and located at the two sides of the D1 domain, respectively (Fig. 6e, h and Fig. S10b). The longer Dc domains of the FlgE subunits of *V. alginolyticus* generate more extensive interactions between the Dc and D1 domains of adjacent subunits of FlgG or FlgE at the -5-, -11-, and -16-start positions in the middle tube (Fig. 6j, k and Fig. S10c), thereby enhancing the rigidity of the hook in the polar motor and generating a tighter tubular junction between the rod and the hook. In addition, the hinge regions (hinge 1 and 2), which connect the D1 and D2 domains, in FlgE of *V. alginolyticus* adopt the conformations that are different from those in the *Salmonella* FlgE (Fig. S11). The D2 domain of FlgE of *V. alginolyticus* contains two distinctive long loops, loop_β5-β6_ and loop_β7-β8_ (Fig. S10d).

**Figure 6.**
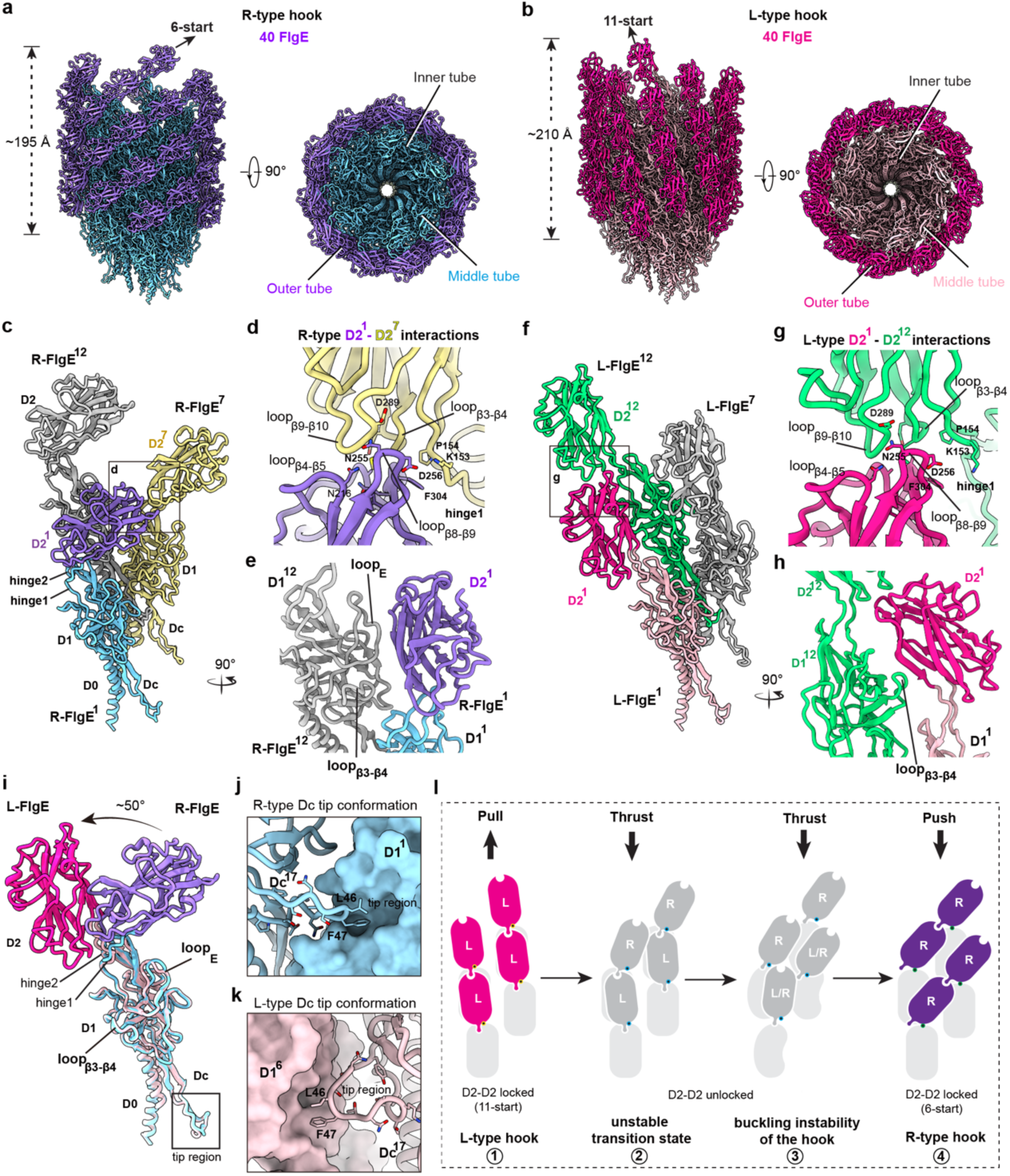
Two types of the hook. (a) Structure of the R-type hook. (b) Structure of the L-type hook. (c) Interactions of FlgE with adjacent FlgE subunits in the R-type hook. (d) Detailed interactions of the D2 domains of the FlgE subunits along the 6-start direction in the R-type hook. (e) Detailed interactions of the D2 domain with the D1 domains of the FlgE subunits along the 11-start direction in the R-type hook. (f) Interactions of FlgE with adjacent FlgE subunits in the L-type hook. (g) Detailed interactions of the D2 domains of the FlgE subunits along the 6-start direction in the L-type hook. (h) Detailed interactions of the D2 domain with the D1 domains of the FlgE subunits along the 11-start direction in the L-type hook. (i) Structural comparison of the FlgE subunit (L-FlgE) in the L-type hook with that (R-FlgE) in the R-type hook. The conformational change of the D2 domain is highlighted with a black arrow. The loop regions of the Dc domains (Dc-loop) are labeled with black lines. (j and k) Comparisons of the detailed interactions of the Dc-loop of FlgE (j) with those in the L-type hook (k). (l) Schematic diagram of a proposed mechanism for the bending and buckling instability of the hook in the “run-reverse-flick” motility pattern.

Structural comparison of the two types of the hook revealed that in the R-type hook, the head of the D2 domain of R-FlgE interacts with the loops at the tail of the D2 domain and the hinge1 and 2 regions of the next R-FlgE subunit along the 6-start direction (Fig. 6d and Fig. S10e). The loop_E_ of the D1 domain at the 11-start position binds to the upper surface of the D2 domain to stabilize the D2-domain protomers in the outer tube of the R-type hook (Fig. 6e). The loop_β3-β4_ regions have no interactions with the D2 domains (Fig. 6e). In the L-type hook, although the D2 domains are packed via the same surfaces that are used in the R-type hook (Fig. S10f), the D2 domains have no interactions with the hinge1 and 2 regions of the L-FlgE subunits along the 6-start direction (Fig. 6g). Instead of loop_E_, loop_β3-β4_ of the D1 domain at the 11-start position binds to the surface of the D2 domain to lock the conformation of the D2-domain protomers (Fig. 6h), suggesting that loop_E_ and loop_β3-β4_ functions as anchors for the outer tube of the L- and R-type hooks, respectively. The tip region of the Dc domain of R-FlgE interact with the D1 domain at the -16-start position (Fig. 6i, j), while the tip region of L-FlgE rotates to interact with the D1 domain at the -5-start position (Fig. 6k). The conformational changes of the tip region of the Dc domain suggest that switching from the L-type to R-type hook can induce the transient disruption of the tight D1-D1 and Dc-D1 interactions in the inner and middle tubes and provide structural flexibility to allow the bending of the hook. In addition, relative to that in L-FlgE, the D2 domain of R-FlgE undergoes a rotation by ∼ 50° via the hinge loops (Fig. 6i and Fig. S10g), which shortens the distances between the D2 domain protomers in the outer tube of the R-type hook and can avoid the structural collapse of the D2 domains in the innermost surface during the bending of the hook.

## Discussion

Bacterial flagellar motors have high diversities in structures for motility in various environments. This study reveals that the polar flagellar motor of *V. alginolyticus* has evolved many distinctive structural elements for high-torque transmission. The additional HT ring enhances the stability of the LP ring, which exerts stronger electrostatic interactions and less physical contacts with the rod to facilitate its high-speed rotation and high-torque transmission. During our submission, an *in situ* cryo-EM structural study reported the additional disk structures, including FlgP- and FlgL-rings, around the LP ring and the MS ring in the polar flagellar motor of *C*. *jejuni* (*38*). All these results highlight the key roles of the structural adaption around the LP ring in the bacterial polar flagellar motors. The unique structure of FlgG and incorporation of phospholipids significantly enhance the rigidity of the rod. In contrast to the 11 rod-binding FliF peptides in the peritrichous flagellar motor (*18*), the MS ring of the polar flagellar motor extends 15 FliF peptide loops to bind the rod for torque transmission.

In the polar flagellar motor, there is also a new component, FlrP, whih has a classic LRR structure with positive charges on surface. The five FlrP subunits binds to the rod and the extended FliF peptide loops, and are located at the space between the P ring and the β-collar region of the MS ring. The protein size and positively-charged surface of FlrP are well complementary to the space and the lower negatively-charged surface of the P ring, respectively, which renders the function of FlrP as the spacers to stabilize the position of the LPHT ring. Consistently, it was observed that the LPHT ring indeed could have slight sliding movements along the rod (*21*). The FlrP subunits can provide a cushioning action for the MS and the LPHT rings against the thrust or pulling force when the bacterial cell swims forward or undergoes the flicking process for the cell reorientation.

There are two types of the hook in the polar flagellum. The packing manners of the D2 domains of the FlgE subunits in the R-type hook can avoid the structural collapse of the D2 domains in the innermost surface during the bending of the hook. Thus, it is possible that the bacterial flicking process requires the state change of the hook from L-type to R-type (Fig. 6l). Upon the motor in the CW directional rotation, the pulling force from the filament pulls the bacterial cell backward and generates the L-type hook by the binding of the loop_β3-β4_ to the D2 domain and the D2-D2 interactions (Fig. 6l). Upon the switching of the rotational direction of the motor to CCW, the thrust force from the filament likely induces the disruption of the D2-D2 and D2- loop_β3-β4_ interactions in the L-hook and reduces the tip-mediated D1-D1 and Dc-D1 interactions to allow the bending of the hook for the bacterial cell reorientation. The D2 domains of FlgE subunits undergoes a rotation by ∼ 50° via the hinge loops to avoid the structural collapses in the innermost side of the hook. After binding by loop_E_ and hinge 1 and 2 of the D1 domains, the D2 domains likely re-establishes the D2-D2 interactions and assemble the R-type hook to push the forward swimming of the bacteria (Fig. 6l). In contrast to the *Salmonella* FlgE, FlgE of *V. alginolyticus* has evolved many structural features, which render the hook not only to be a joint to connect the filament, but also to be a rudder for bacterial cell reorientation in swimming.

## Materials and Methods

### Bacterial strains and culture conditions

The *Vibrio alginolyticus* strains VIO5(*39*) and KK148(*33, 34*) were cultured at 30°C in VC medium consisting of 0.5% (w/v) tryptone, 0.5% (w/v) yeast extract, 3% (w/v) NaCl, 0.4% (w/v) K_2_HPO_4_, and 0.2% (w/v) glucose. *Escherichia coli* stains DH5α λπ and SM10 λπ were grown at 37°C in 2YT medium. Chloramphenicol and ampicillin were added at the final concentrations of 25 μg/mL and 10 μg/mL, respectively.

### Construction of *V. alginolyticus* mutant strains

The *V. alginolyticus* KK148 strain is a multipolar flagellar mutant strain derived from the VIO5 strain with a *flhG* gene mutation, producing several polar flagella at one cell pole(*33*). To eliminate the flagellar filament contamination during purification, the *flgL* gene, which encodes hook-associated protein 3 (HAP3) for connecting the filament to the hook, was deleted by using a homologous recombination-based gene knockout system (*14, 40*). The 1000 bp sequences upstream and downstream of the *flgL* coding region from the KK148 strain were amplified as the homologous arms to construct the pDM4-*ΔflgL* suicide plasmid. This plasmid includes a chloramphenicol resistance gene and an R6K replication origin dependent on the π protein. To avoid disrupting adjacent genes, 60 bp on each side of the *flgL* gene were retained. The obtained pDM4-*ΔflgL* plasmid was transformed into the *E. coli* SM10 λπ donor strain. The suicide plasmid-containing SM10 λπ strain and the recipient KK148 strain were separately cultured in antibiotic-free 2YT and VC medium, respectively, for six hours. The suspension cultures were mixed and deposited onto the 0.22 μm filter membrane placed on VC plate and incubated overnight at 30°C for conjugation. The mated culture was diluted and spread on VC plates containing 10 μg/mL chloramphenicol and 100 μg/mL ampicillin, then incubated overnight at 30°C for the first-round recombination. Single colonies were then grown in VC medium at 30°C and identified using primers specific to the *sacB* gene. The *sacB*-positive clones were streaked onto the sucrose plates containing 10% sucrose, 1% tryptone, 30 mM NaCl, 55 mM KCl and 1.25% agar, and incubated overnight at 30°C for the second-round recombination. The obtained mutant stain KK148-*ΔflgL* was finally validated by PCR and sequencing. The *V. alginolyticus* VIO5 strain is a lateral flagella-deficient strain with a single polar flagellum at the cell pole. The VIO5-*ΔflrP* strain was constructed using the same methods described for the construction of the KK148-*ΔflgL* stain. To construct the *V. alginolyticus* VIO5-Δ*flrP::flrP* mutant strain, the *flrP* gene was complemented into the genome of VIO5-*ΔflrP* strain using the pDM4 suicide vector.

### Purification of the polar motor-hook complex

The KK148-*ΔflgL* strain was cultured overnight at 30°C in 5 mL of VC medium. The 1.6 mL of the culture was inoculated into 1 L of VC medium supplemented with 0.1% glucose and grown at 30°C for 4.5 h. The cells were harvested by centrifugation at 4,000 rpm and resuspended in 100 mL of lysis buffer consisting of 100 mM Tris-HCl (pH 8.0), 0.5 M sucrose, 10 mM EDTA (pH 7.5), and 0.1 mg/mL lysozyme. The suspension was stirred on ice at 200 rpm for 1 h. The cells were then lysed by adding Triton X-100 to a final concentration of 1% (v/v) and 20 mM MgSO_4_ at room temperature. To digest endogenous nucleic acids and clarify the cell lysates, 200 mg DNase I powder (BBI Life Sciences Corporation) was added to the mixture and incubated at room temperature for 10 minutes. The additional 100 mL of lysis buffer pre-mixed with 1% (v/v) Triton X-100 was then added, followed by incubation at room temperature for 10 minutes and at 4°C for 1 h. The cell debris was removed by centrifugation at 14,600 rpm for 20 min. The pH value of the supernatant was adjusted to 10.5 using 5 M NaOH, and denatured proteins were then removed by centrifugation at 14,600 rpm for 15 min. Subsequently, the supernatant was centrifugated at 35,000 rpm for 1 h at 4°C to obtain the membrane pellet containing the bacterial polar flagellar motor-hook complex particles. The pellet was resuspended in 3 mL of TET buffer (10 mM Tris-Cl, pH 8.0; 5 mM EDTA, pH 7.5; 0.1% Triton X-100) and further homogenized using a Dounce homogenizer (7 mL Tissue Grinder, Dounce, #357542). The suspension was incubated at 4°C for 1 h and clarified by centrifugation at 12,000 rpm for 10 min. The supernatant was then loaded onto a 20%-50% sucrose gradient in SW41 tubes and centrifuged at 20,000 rpm (∼68,000 g) for 12 h at 4°C. The fractions containing the bacterial flagellar motor-hook complex particles were collected from the bottom of the tube in 200 μL aliquots. These fractions were analyzed by negative staining, and those containing the bacterial polar flagellar motor-hook complex were then pelleted by centrifugation at 20,000 rpm for 2 h. The transparent pellet was resuspended in 10 μL of TET buffer and immediately used for subsequent cryo-EM analyses for the motor-hook complex. The purification method of the LPHT ring was identical to that for the motor-hook complex, except that the double-distilled water, bot the TET buffer, was used to prepare the sucrose gradient in centrifugation for isolating the disassociated LPHT ring particles.

### Negative staining analysis

The 2.5 μL of the samples were loaded to a glow-discharged carbon-coated copper grid (200 mesh) for 2 min at room temperature, after which the solution was removed using filter paper. The grid was stained with 5 μL of 2% (w/v) uranyl acetate for three times, 10 s for the first and second staining, and 1 min for the third staining. The grids were air-dried at room temperature after removing the uranyl acetate. Grids were visualized using a transmission electron microscope (HT7700, HITACHI) operated at 80 kV.

### Cryo-EM sample preparation

The 3 μL of the samples were applied onto the ultra-thin carbon film supported grids (QUANTIFOIL R1.2/1.3 300 Cu, 2nm) and then plunge-frozen using an FEI Vitrobot Mark IV. Before use, the grids were glow-discharged by PELCO easiGlow with the condition of 10 mA for 10 s. Blotting was performed for 3 s with a blotting force of 15, followed by a 60-second wait time and 2-second drain time at 100% humidity and 4°C. These parameters were applied for the sample preparation of the motor-hook complex and the LPHT ring. All these cryo-EM grids were then stored in liquid nitrogen for data collection.

### Cryo-EM data collection

Data collection was conducted using FEI Titan Krios electron microscope operated at voltage of 300 kV equipped with a Falcon 4 camera. For the cryo-EM images of the motor-hook complex that mainly contain the end-on views of the disassociated LPHT ring particles, images were recorded using the software of EPU (Thermo Fisher Scientific) at a nominal magnification of 105,000. A total of 3,600 electron-event representation (EER) movies were acquired with defocus value ranging from 1.2 to 1.8 μm. The calibrated pixel size was set to 1.2 Å/pixel and the exposure time was set to 6.86 seconds, resulting in a total exposure dose of ∼50 electrons/Å^2^. For the cryo-EM images of the LPHT ring that mainly contain the particles of the intact polar flagellar motor-hook complex, a total of 11,887 EER movies were captured using the same parameters.

### Cryo-EM Image processing

The total of EER movies were imported into the CryoSPARC software(*41*), followed by the motion correction and contrast transfer function (CTF) estimation. For the cryo-EM images of the motor-hook complex (3,684 micrographs), 3,284 particles of the LPHT ring in end-on views were obtained using template-based auto-picking with a box size of 560×560 pixels. Preliminary 2D averages revealed the 26-fold symmetry of the LPHT ring. For the cryo-EM images of the LPHT ring (11,887 micrographs), which contained the particles of the intact polar flagellar motor-hook complex, a total of 329,955 particles centered on the LPHT ring were automatically picked with a box size of 620×620 pixels. After multiple rounds of 2D classification, 61,395 particles were selected for ab-initio three-dimensional (3D) reconstruction. Subsequent homogeneous refinement applied with C26 symmetry produced a 2.8-Å density map of the LPHT ring. Due to the mixed conformations and the poor quality of the T ring at the bottom of the LPHT ring, we subtracted the signal of the central axial structure with the MS ring, and performed unsupervised 3D classification (without alignment, 4 classes) on the T ring region after non-uniform refinement (C26). Two main classes revealed that T ring has 13-fold symmetry, while the LPH ring has 26-fold symmetry. A total of 45,233 particles (73.8%) were subjected to the non-uniform refinement applied with C13 symmetry, yielding a 2.95-Å density map of the LPHT ring. We then removed the signal of the LPHT ring, which affected the overall structure calculation from 61,395 particles. Another round of 2D classification yielded 58,419 particles for ab-initio 3D reconstruction (C1) and non-uniform refinement (C1), generating a 3.5-Å density map of the region containing the hook, the rod, the export apparatus and the MS ring. Further CTF and local refinements produced a 3.0-Å density map of the region containing the export apparatus, the rod and a part of the hook (FlgE^1-11^-D0-D1 domains), and an additional local refinement produced a 3.0-Å density map of the proximal rod with the export apparatus. We then re-centered particles on the MS ring and produced a 3.6-Å density map of the MS ring region containing the MS ring, the export apparatus and a part of the proximal rod after homogeneous and local refinements with C1 symmetry. Further local refinement of the β-collar-RBM3 subrings with C34 symmetry improved its resolution to 2.9 Å.

A 3D classification (without alignment, 4 classes) focusing on the proximal rod region was performed due to the five extra densities observed near the proximal rod and the MS ring in the 3.5-Å density map of the region containg the hook, the rod, the export apparatus and the MS ring. Further CTF and local refinements of 19,450 particles (33.3%) in one class, featuring two β-solenoid-shaped densities, yielded a middle rod density map at 3.5 Å, with two β-solenoid structures bound to the outer surface of the proximal rod. Subsequently, all particles were re-centered on the hook region and reextracted with a box size of 380×380 pixel. After 2D classification yielding 57,802 particles, two distinct conformations of the hook were identified: 34,667 particles of the L-type hook and 20,187 particles of the R-type hook. Ab initio reconstruction and nun-uniform refinement with C1 symmetry were performed, followed by 3D classification (without alignment, 4 classes) and helical refinements, generating a 3.2-Å density map of the L-type hook and a 3.4-Å density map of the R-type hook. Finally, a total of 61,395 particles were subjected to homogeneous refinement applied with C100 symmetry and produced the overall density map of the polar flagellar motor-hook complex, which served as the reference coordinate for the composite density map and model building.

All map sharpening was performed automatically by CryoSPARC, which also generated the Fourier Shell Correlation (FSC) curves for the reconstructions. Average resolutions were estimated from the corrected FSC curves at the 0.143 threshold. Local resolution maps were calculated in CryoSPARC using the two independent half-maps from each reconstruction as input and visualized with *ChimeraX*(*42*). All volume masks for local refinements were created using *Chimera*(*43*).

### Structure prediction by AlphaFold2

Structure predictions of FlrP for model building were calculated using ColabFold (https://github.com/sokrypton/ColabFold) plugged in *ChimeraX* (*44*). A total of five models were generated for each prediction, and the best model was selected as an initial model for model building. Predicted structures were visualized and colored in *ChimeraX* based on the predicted local distance difference test (pLDDT) scores. The predicted aligned error (PAE) plot, which indicates the expected positional error of the predicted complex, was automatically generated by AlphaFold2 (*45*).

### Model building and validation

The structures of the FlgH, FlgI, FlgT, FlgPc, MotX and MotY monomers were manually modeled based on the initial model generated by *ModelAngelo* (*46*) and refined in the 2.95-Å density map of the LPHT ring (C13), and produced the protomers of the LPH ring (FlgH_1_-FlgI_1_-FlgT_1_-FlgPc_1_) and the T ring (MotX_1_-MotY_1_, MotX_2_-MotY_2_). After multiple rounds of real-space refinement in *Phenix* (*47*) with manual inspection and adjustments in *Coot* (*48*), the final models of the LPH ring and the T ring were generated in *ChimeraX* using C26 and C13 symmetry, respectively. The overall structure of the LPHT ring was obtained by combining the LPH ring and T ring after rigid-body refinement in *Phenix*. The structures of FliE, FlgF, FlgC, FlgB, FlgG, FliP, FliQ, FliR, FlhB, and FlgE-D0-D1 monomers were manually built and refined in the 3.0-Å density map of the region contain the rod, the export apparatus and a part of the hook and the map of the proximal rod with the export apparatus in *Coot* and *Phenix*, respectively, based on the initial models generated by *ModelAngelo*. The 6 FliE, 5 FlgB, 6 FlgC, 5 FlgF, 27 FlgG, 5 FliP, 4 FliQ, 1 FliR, 1 FlhB and 11 FlgE-D0-D1 subunits were generated according to the helical symmetry in *ChimeraX*. The models of the proximal rod with the export apparatus and of the distal rod with a part of the hook were obtained after several iterations of manual adjustments in *Coot* and real-space refinement in *Phenix* within the corresponding density maps. The final overall model of the rod-export apparatus-partial hook was obtained by combining the above two refined models and imported into *Phenix* for an additional round of rigid-body refinement.

The protomer of the β-collar-RBM3 subrings was built and refined in the 2.9-Å density map of the MS ring (C34), while the protomer of the RBM2 subring was built and refined in the 3.6-Å density map of the MS ring with the export apparatus and a part of the proximal rod (C1), based on the initial models generated by AlphaFold2. The overall model of the β-collar-RBM3 subrings was obtained using C34 symmetry in *ChimeraX* and refined with NCS (non-crystallographic symmetry) restraints in *Phenix*. The model of the RBM2 subring (C23) were adjusted individually in *Coot* and refined without NCS restraints in *Phenix*. The final model of the MS ring-export apparatus-proximal rod was obtained by combining the refined models of the β-collar-RBM3 subrings, the RBM2 subring, the export apparatus and the part of the proximal rod, and then imported into *Phenix* for a round of rigid-body refinement in the 3.6-Å density map of the MS ring (C1). The models of the FliF-L_A_^1-5^, FliF-L_B_^1-5^, and FliFL-_C_^1-5^ peptide loops were built in the 3.0-Å density map of the proximal rod with the export apparatus and the 3.5-Å density map of the middle region of the rod, based on the initial model generated by *ModelAngelo*. The FlrP^1^ and FlrP^5^ models were built in the 3.5-Å density map of the middle region of the rod based on the initial model generated from *ModelAngelo* and AlphaFold2. The final models were adjusted in *Coot* and then refined using *Phenix*.

The model of the R-FlgE subunit in the R-type hook was manually adjusted in *Coot* and further refined in *Phenix* in the 3.4-Å density map of the R-type hook, based on the initial model generated by AlphaFold2. The initial model of the L-FlgE subunit in the L-type hook was derived from the R-FlgE subunit and refined using *Phenix* in the 3.2-Å density map of the L-type hook. The models of the L-type hook and R-type hook were generated using the helical symmetry command in *ChimeraX*, incorporating 40 L-FlgE and 40 R-FlgE subunits in their corresponding density maps, and individually adjusted in *Coot* and refined using *Phenix*. The model of the L-hook-rod-export apparatus was obtained by aligning the L-hook with the rod-export apparatus-partial hook and FlrP in the 3.0-Å density map of the region containing the rod, the export apparatus and the part of the hook. The MS ring, the export apparatus, and the part of the proximal rod were also modeled in this density map to generate the L-hook-rod-export apparatus-MS ring model. This model with hat of the LPHT ring was then fitted into the density map of the overall polar flagellar motor-hook complex (C100) using *ChimeraX*. The final overall model of the polar flagellar motor-hook complex was composited by combining these two models and validated using *MolProbity* without further refinements.

All structural models were validated using *Molprobity* (*44*). The statistical data of data collection, processing, model refinement and validation are listed in Table S1. Representative structures and electron density are shown in Fig. S2B. Structural superpositions were performed using the in *ChimeraX*. The electrostatic distribution was visualized in *ChimeraX.* All Figures were generated using *ChimeraX*.

### Mass spectrometric analysis

The SDS-PAGE gel containing the protein sample band was cut and digested by trypsin at 37°C overnight with prior reduction and alkylation in 50 mM ammonium bicarbonate. The extracted peptides were then subjected to NSI source followed by tandem mass spectrometry (MS/MS) in Q ExactiveTM Plus (Thermo) coupled online to the UPLC. The resulting MS/MS data were processed using Proteome Discoverer 1.3. Tandem mass spectra were searched against NCBI database (GenBank: AP022861.1, GenBank: AP022862.1). Mass error was set to 10 ppm for precursor ions and 0.02 Da for fragmental ions. Carbamidomethyl on cysteine were specified as fixed modification and oxidation on Met was specified as variable modification. Peptide confidence was set at high, and peptide ion score was set > 20.

### Bacterial motility assay

To compare the motility of the VIO5 strain, the VIO5-*ΔflrP* strain and the complementary VIO5-*ΔflrP::flrP* strain, all strains were inoculated in 5 mL of antibiotic-free VC medium and cultured overnight. A 1 μL aliquot of the overnight cultures was spotted onto the VPG soft agar plate containing 0.25% (w/v) agar, 1% (w/v) tryptone, 0.4% (w/v) K_2_HPO_4_, 3% (w/v) NaCl, and 0.5% (w/v) glycerol. The volume of the remaining strains was adjusted, based on their OD_600_ values. The plates were incubated at 30°C for 6 h, and the diameters of the swimming circles were measured using ImageJ. This experiment was repeated 10 times, and the data were analyzed using GraphPad Prism 9.0 software.

### Western blot

To determine the presence of the FlrP protein, the western blot was performed using the purified polar flagellar motor-hook complex particles. The samples were mixed with SDS loading buffer and boiled at 95 °C for 5 min, then applied to a 15% SDS-PAGE gel. After electrophoresis, the proteins were transferred onto PVDF membranes. The custom-made FlrP monoclonal antibody (rabbit source, HUABIO, #HA722715) was diluted in the TBST buffer (1:5000) and incubated for 1 h at room temperature. The basic ECL light-emitting substrate was used for chemiluminescence after a 1-hour incubation at room temperature with goat anti-rabbit IgG conjugated to horseradish peroxidase (1:10000). The PVDF membrane was then stained with Coomassie blue to visualize the bands.

### Quantification and statistical analysis

In Figure S1 and S2, the tight and corrected FSC curves of each reconstruction were calculated using a tight mask alone and a tight mask with noise substitution correction based on two independent half-maps in CryoSPARC. These curves were illustrated using GraphPad Prism 9.0. The resolution estimations of the cryo-EM density maps were calculated in CryoSPARC using the two independent half-maps of each reconstruction as input, based on the corrected FSC curves at the FSC=0.143 criterion. The model resolution in Table S1 was evaluated using *phenix.mtriage* from the model-based noise free map and the experimental map at the FSC=0.5 criterion. Angle and distance measurements were performed in *ChimeraX*.

## Acknowledgments

We thank the core facility of Life Sciences Institute Zhejiang University for equipment support and the cryo-EM centers of Zhejiang University for their assistance in cryo-EM data collection. This work was supported by grants from NSFC (U23A20163 and 82525106 to Y. Zhu, 82172282 to Y. Zhou), the National Key R&D Program of China (2024YFA1306800), and the Fundamental Research Funds for the Central Universities. Y. Zhou is supported by the National High-level Talents Special Support Program. Y.

Zhu is supported by New Cornerstone Science Foundation.

## Author contributions

Y. Zhu and Y. Zhou conceived and supervised the study; L.Z., J.T. and Y. Zhu designed experiments; M. H. and S. K. provided the KK148 strain; L.Z. and J.T. purified the complexes and determined cryo-EM structures; X.D. and X. W. assisted assays; L.Z., J. T., Y. Zhou and Y. Zhu prepared the manuscript.

## Data and materials availability

The cryo-EM density maps and structural models of the Vibrio polar flagellar motor-hook complexes have been deposited in the EMDB and PDB with the following accession numbers: the LPHT ring structure, EMDB: EMD-39776, PDB: 8Z5N; the Rod-Export apparatus partial hook complex, EMDB: EMD-39782, PDB: 8Z5U; the Proximal rod-Export apparatus, EMDB: EMD-39780, PDB: 8Z5S; the partial Rod, EMDB: EMD-39786, PDB: 8Z5W; the β-collar-RBM3 subrings of the MS ring with C34 symmetry, EMDB: EMD-39783, PDB: 8Z5V; the MS ring (C1, containing partial proximal rod and export apparatus), EMDB: EMD-39787, PDB: 8Z5X; the L-type hook, EMDB: EMD-39788, PDB: 8Z5Y; the R-type hook, EMDB: EMD-39789, PDB: 8Z5Z; and the overall polar flagellar motor-hook complex with C100 symmetry, EMDB: EMD-39790, PDB: 8Z60.

## Supplementary materials

**Figure S1:**
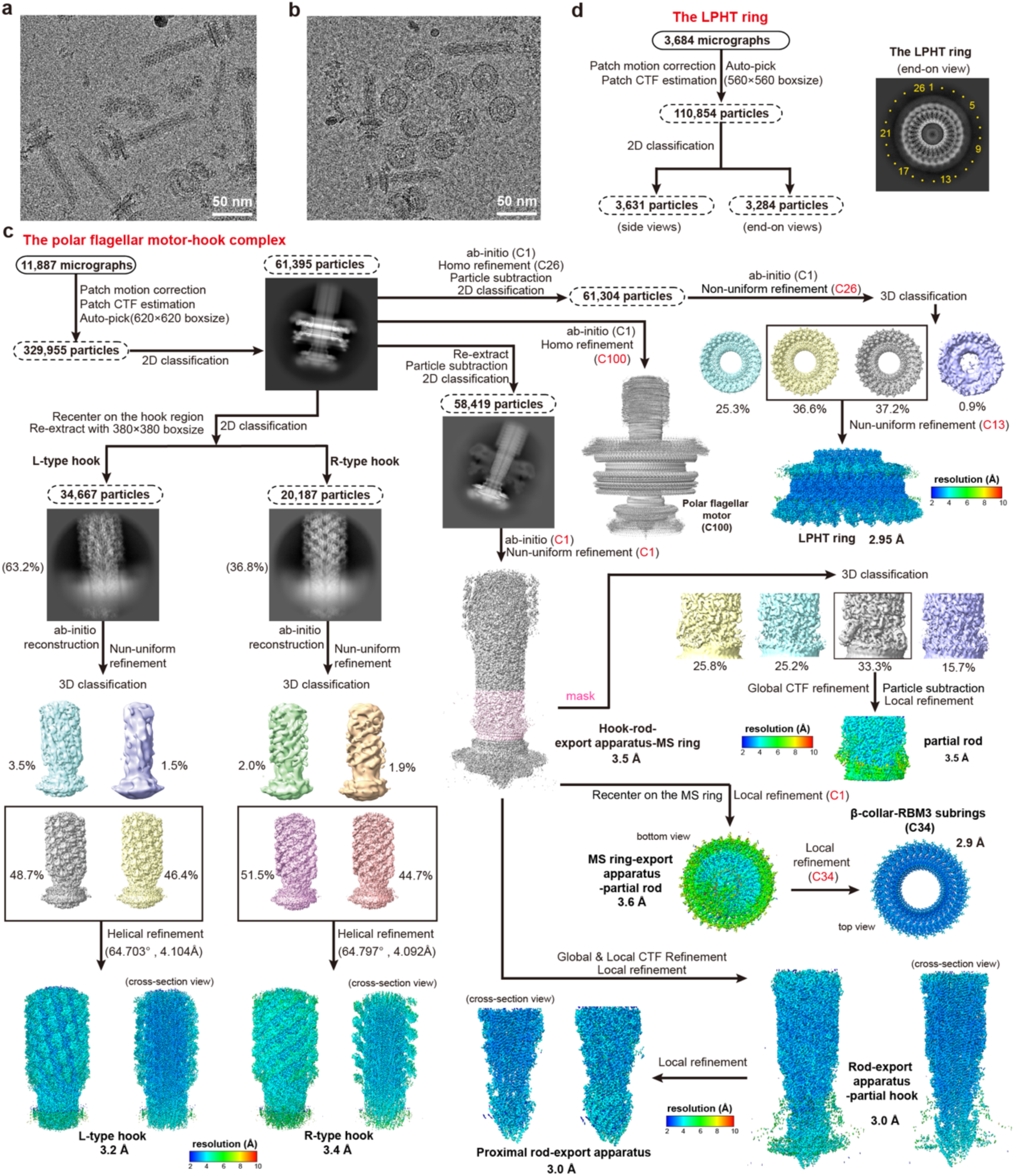
Cryo-EM data processing of the polar flagellar motor-hook complex. (a) A representative cryo-EM micrograph of the polar flagellar motor-hook complex. (b) A representative cryo-EM micrograph of the particles of the LPHT ring collected. Scale bar, 50 nm (a, b). (c) The flow chart of the cryo-EM data processing and structural determination of the polar flagellar motor-hook complex. (d) Flow chart of the cryo-EM data processing of the LPHT ring.

**Figure S2:**
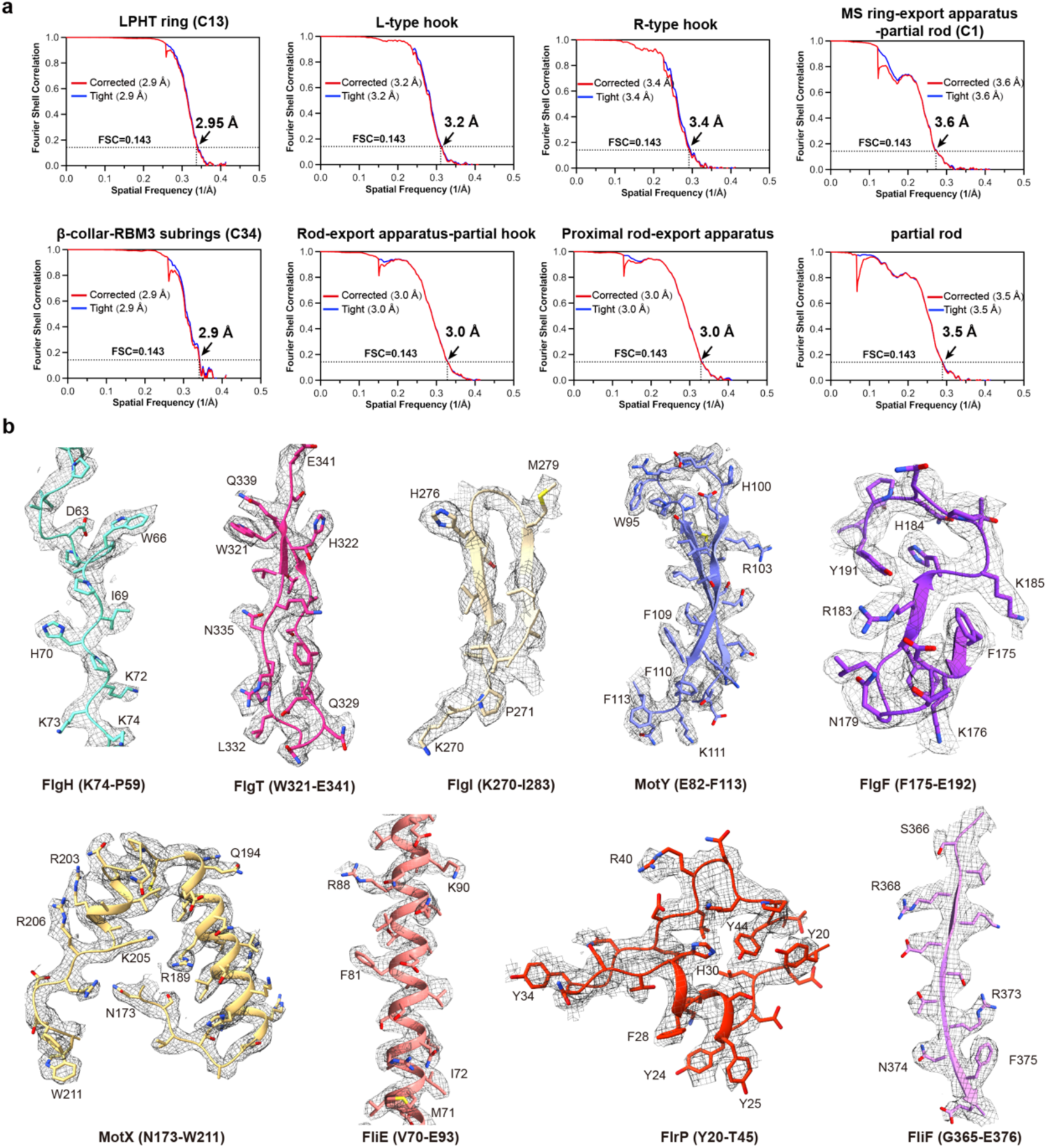
FSC curves and representative cryo-EM density maps of the polar flagellar motor-hook complex. (a) The FSC (Fourier Shell Correlation) curves and estimated resolutions for the reconstructions in (Fig. S1c). (b) Representative cryo-EM density maps of FlgH, FlgT, FlgI, MotY, FlgF, MotX, FliE, FlrP and FliF after local refinements.

**Figure S3:**
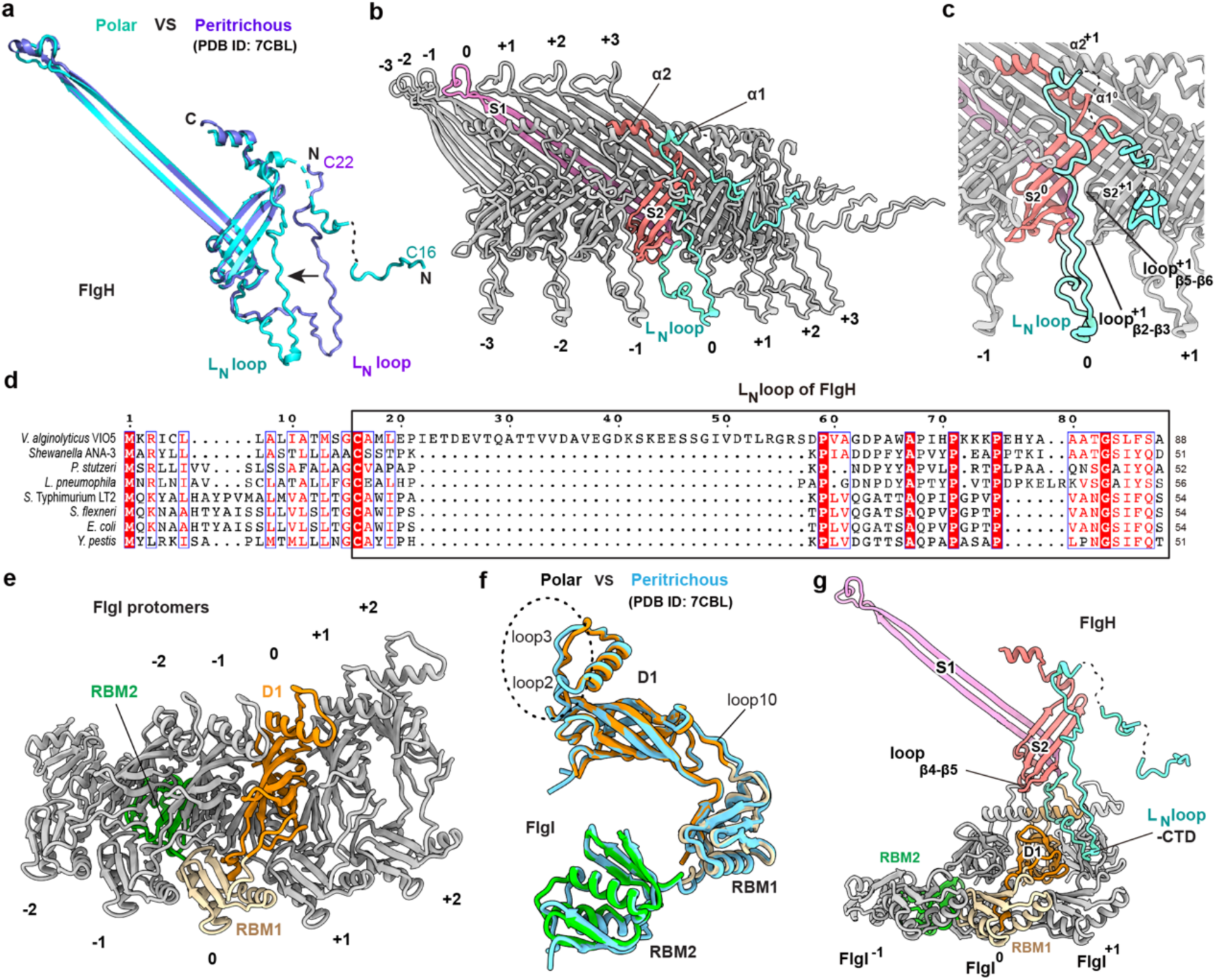
Structures of FlgH and FlgI in the polar LP ring. (a) Structural comparison of FlgH in the polar flagellar motor-hook complex with that in the peritrichous flagellar motor-hook complex. The polar and peritrichous FlgHs are colored in cyan and blue, respectively. (b) Inter-subunit interactions of FlgH in the L ring in the polar flagellar motor. The S1, S2 β-sheets and L_N_ loop of the polar FlgH are colored in pink, orange and cyan, respectively. (c) Close-up view of the L_N_ loop and its interactions with adjacent FlgH subunits in the L ring. (d) Sequence alignment of the L_N_ loop of FlgH of *V. alginolyticus* with those of other bacterial species. (e) Inter-subunit interactions of FlgI in the P ring. (f) Structural comparison of FlgI in the polar flagellar motor-hook complex and that in the peritrichous flagellar motor-hook complex. The D1, RBM1 and RBM2 domains of the polar FlgI are colored in orange, yellow and green, respectively. The peritrichous FlgI is colored in cyan. (g) The 1-versus-3 interaction manner of the FlgH and FlgI subunits with each other between the L and P rings.

**Figure S4:**
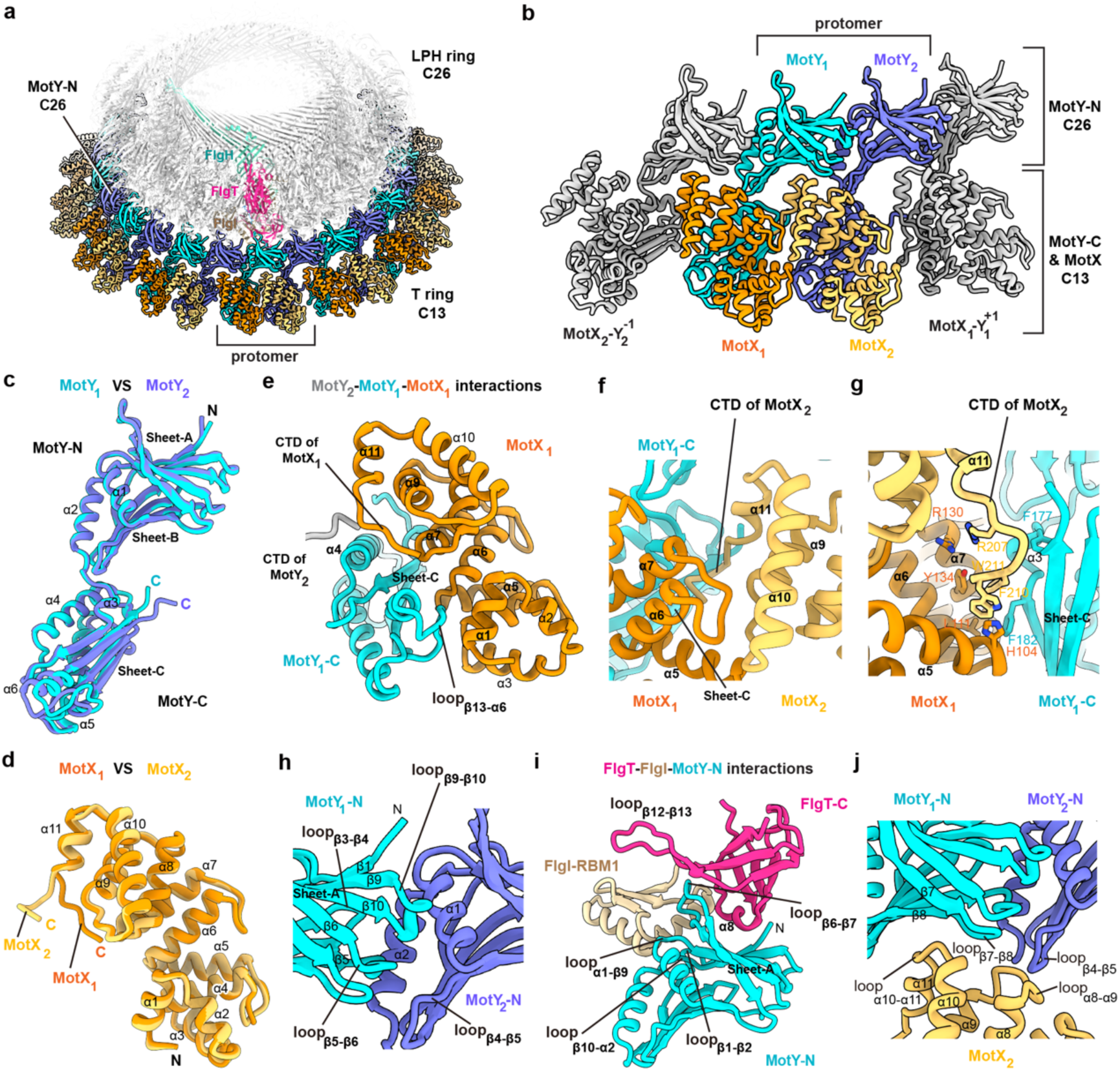
The symmetry mismatch in the LPHT ring structure. (a) Overall view of the structure of the T ring. Each protomer of the T ring consists of MotX_1_, MotY_1_, MotX_2_ and MotY_2_, which are colored in orange, cyan, gold and deep blue, respectively. The LPH ring is colored in grey. The subunits of FlgH, FlgI and FlgT in a protomer of the LPH ring are colored in cyan, yellow and red, respectively. (b) Close-up view of the protomer of the T ring. (c) Structural comparison of the two MotY subunits of the protomer. (d) Structural comparison of the two MotX subunits of a protomer. (e) Interactions between MotX_1_ and MotY_1_ in the protomer. (f and g) Detailed interactions of the CTD region of MotX_2_ with MotX_1_ and MotY_1_-C in a protomer. (h) Interaction of the N-terminal domains of MotY_1_ and MotY_2_ in a protomer. (i) Interactions between the MotY-N domain and the upper LPH ring. (j) Interactions of MotX_2_ with the MotY-N domains in a protomer.

**Figure S5:**
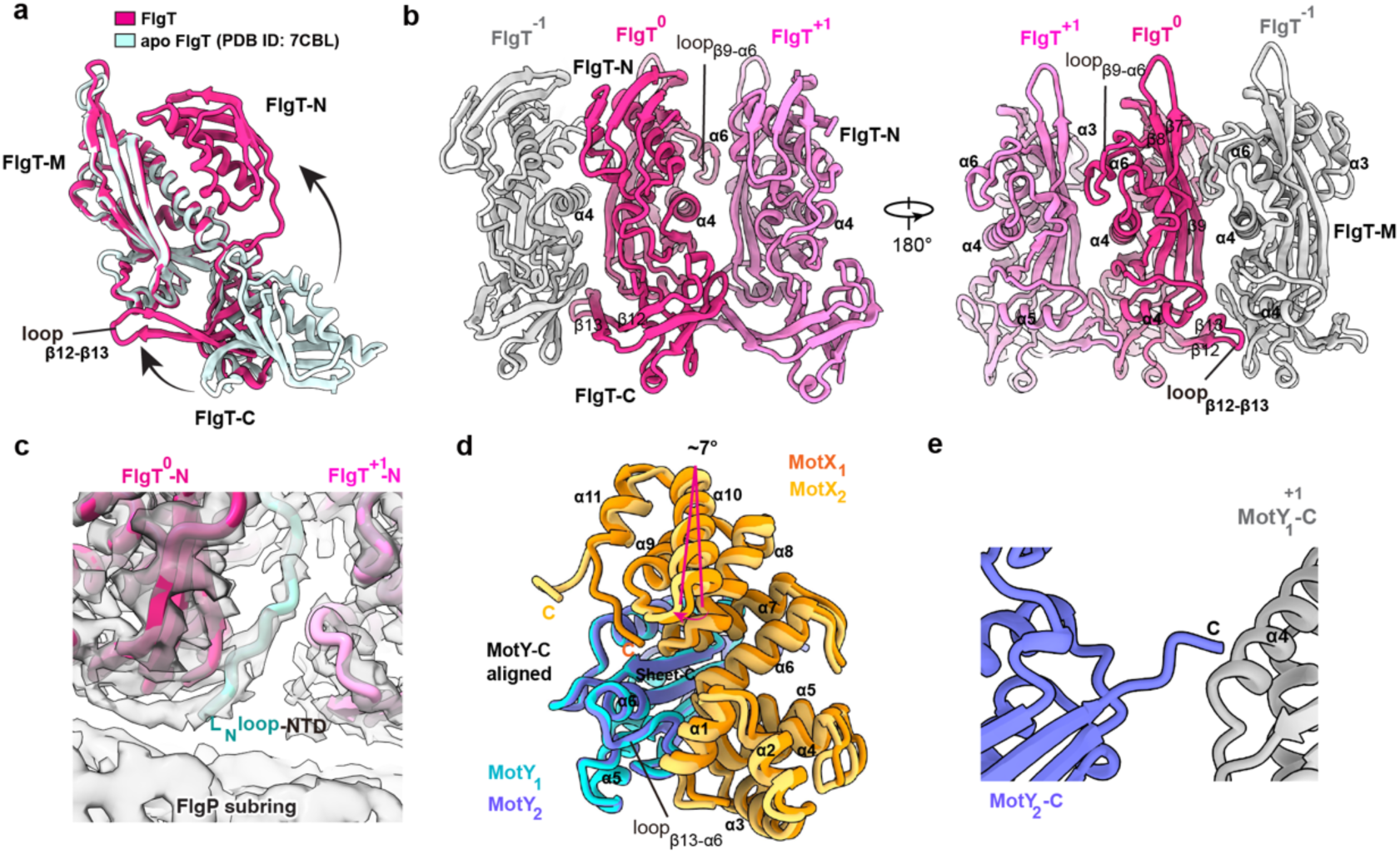
Inter-subunit interactions in the H and T rings. (a) Structural comparison of FlgT in the polar flagellar motor-hook complex with its apo structure (PDB ID: 3W1E). (b) Inter-subunit interactions of the FlgT subunits in the H ring. (c) Interactions of the N-terminal region of the L_N_ loop of FlgH with the H ring. (d) Structural superimposition of the MotX_1_-MotY_1_ and MotX_2_-MotY_2_ dimers of a protomer, based on the C-terminal domains of MotY. (e) The inter-protomer interaction of the C-terminus of MotY_2_ with the MotY_1_ subunit of the next protomer in the T ring.

**Figure S6:**
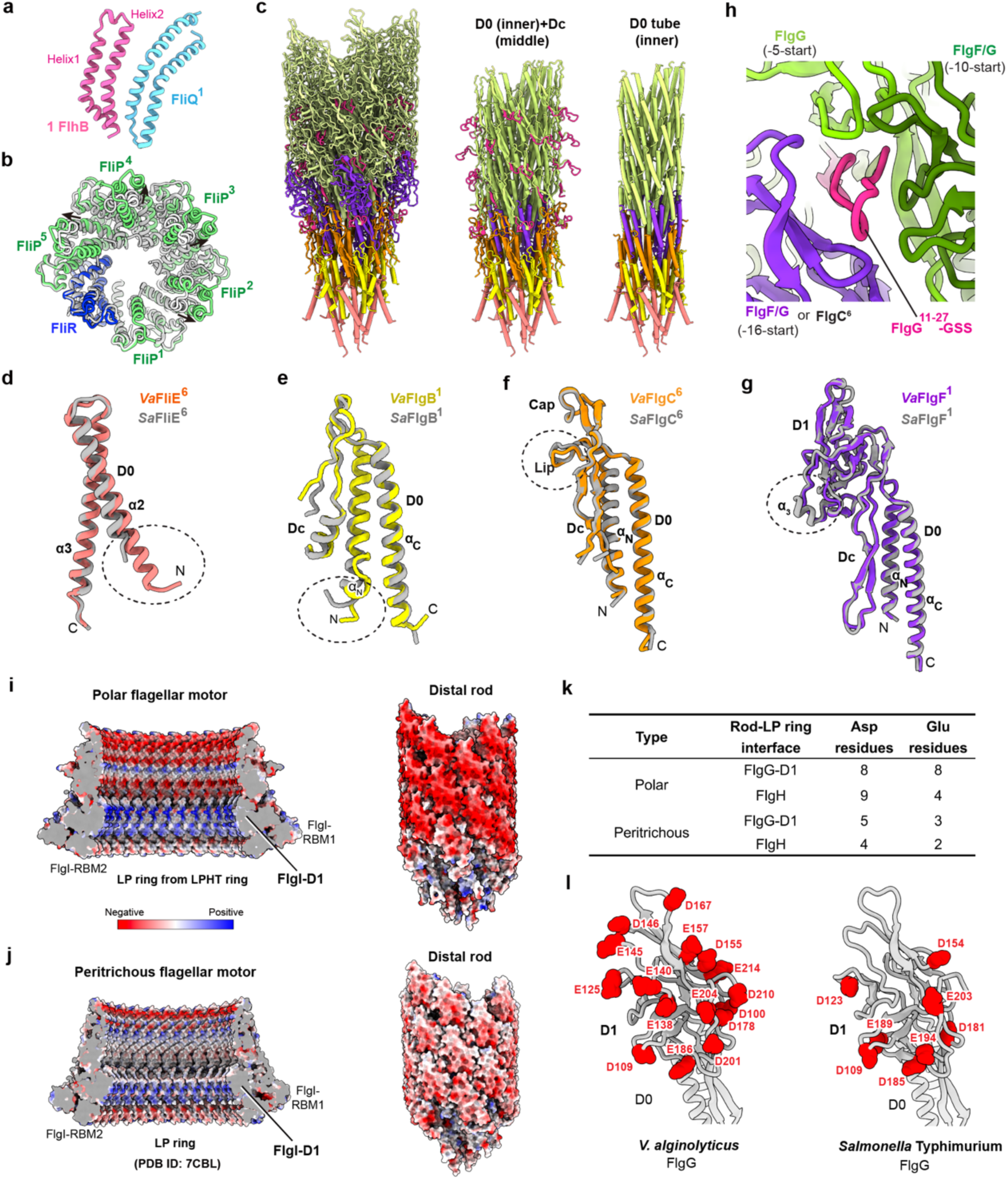
The export apparatus, and the structures of the rod components in the polar flagellar motor. (a) Structural comparison of FlhB and FliQ. (b) Structural comparison of the proximal rod-binding export apparatus in the polar flagellar motor-hook complex with the free export apparatus (PDB ID: 6S3L). The free export apparatus is colored in grey. (c) The three-layered tubular structure of the rod in the polar flagellar motor-hook complex. The subunits of FliE, FlgB, FlgC, FlgF and FlgG are colored in salmon, yellow, orange, purple and light green, respectively. (d-g) Structural comparison of the rod components in the polar and *Salmonella* peritrichous flagellar motor-hook complexes. (h) Interactions of the FlgG^11-27^-GSS region with adjacent subunits in the rod. (i and j) Comparison of the electrostatic potential surfaces of the LP ring (left) and the rod (right) in the polar and peritrichous flagellar motor-hook complexes. (k) Comparison of the numbers of the aspartic acid (Asp, D) and glutamic acid (Glu, E) residues on the surface of FlgH and the D1 domain of FlgG at the rod-LP ring interface. (l) Distribution of the aspartic acid and glutamic acid residues on the surfaces of the D1 domains of FlgGs of *V. alginolyticus* (left) and *S.* Typhimurium (right).

**Figure S7:**
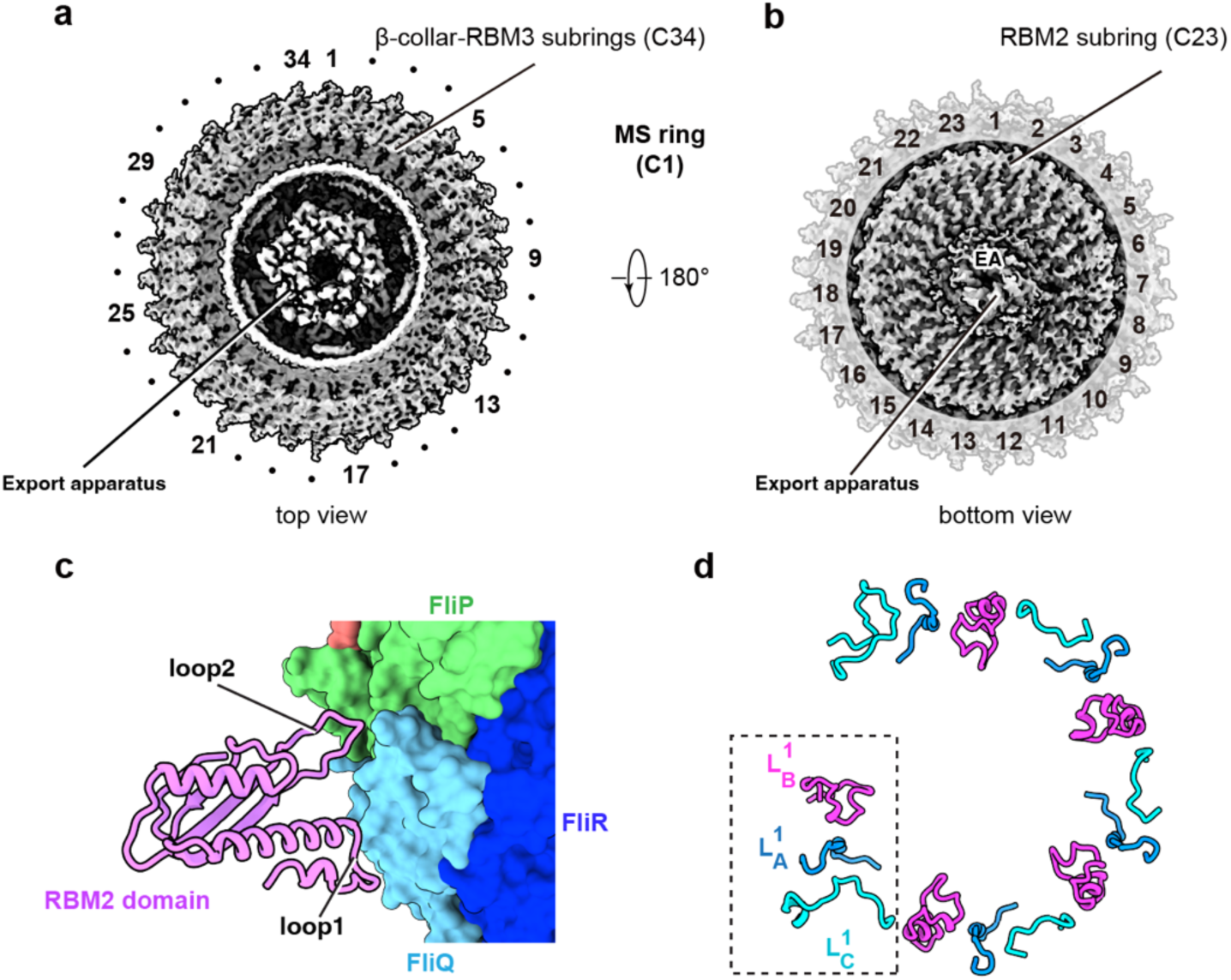
Structure of the MS ring in the polar flagellar motor-hook complex. (a and b) Top (a) and bottom (b) views of cryo-EM density map of the MS ring constructed by C1 symmetry. The β-collar-RBM3 subrings have the C34 symmetry (a), while the RBM2 subring exhibits the C23 symmetry (b). (c) Interaction of the RBM2 domain of FliF with the export apparatus through Loop 1 and 2 in the polar flagellar motor-hook complex. (d) Top view of the distribution of the 15 FliF peptide loops. The loops are grouped into 5 sets, each of which contains a L_A_, L_B_ and L_C_ loops.

**Figure S8:**
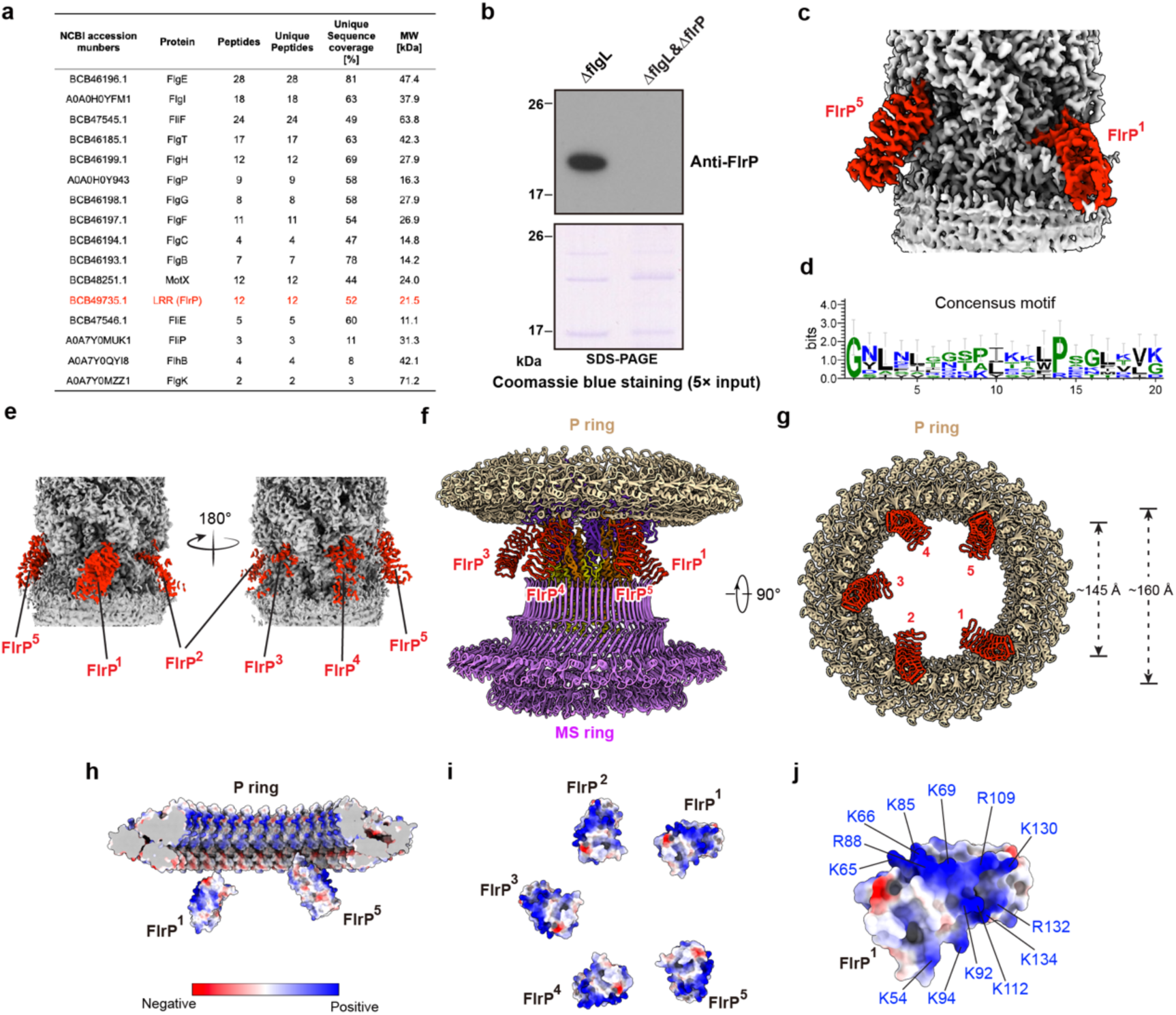
Identification of FlrP in the polar flagellar motor. (a) Mass spectrometry analysis of the components in the purified particles of the polar flagellar motor-hook complex. The leucine-rich repeat (LRR) protein FlrP is highlighted in red in the table. (b) Western blot examination of FlrP in the purified particles of the polar flagellar motor-hook complex. Upper, western blot; blow, SDS-PAGE with Coomassie blue staining. (c) Representative cryo-EM density map of FlrP with the rod. The densities of FlrP^1^ and FlrP^5^ are illustrated and colored in red. (d) The sequence consensus of the leucine-rich repeated coils of FlrP. (e) Structural model of the five FlrP subunits bound to the rod. (f) Bottom views of the positions of the FlrP subunits under the P ring. (g) Cross-section view of the electrostatic potential distributions of the surfaces of the FlrP subunits and the P ring. (h) The electrostatic potential distributions of the five FlrP subunits. (i) The positively charged residues on the surface of FlrP. The surface is colored by relative electrostatic potential with red indicating negatively charged and blue positively charged (g, h, and i).

**Figure S9:**
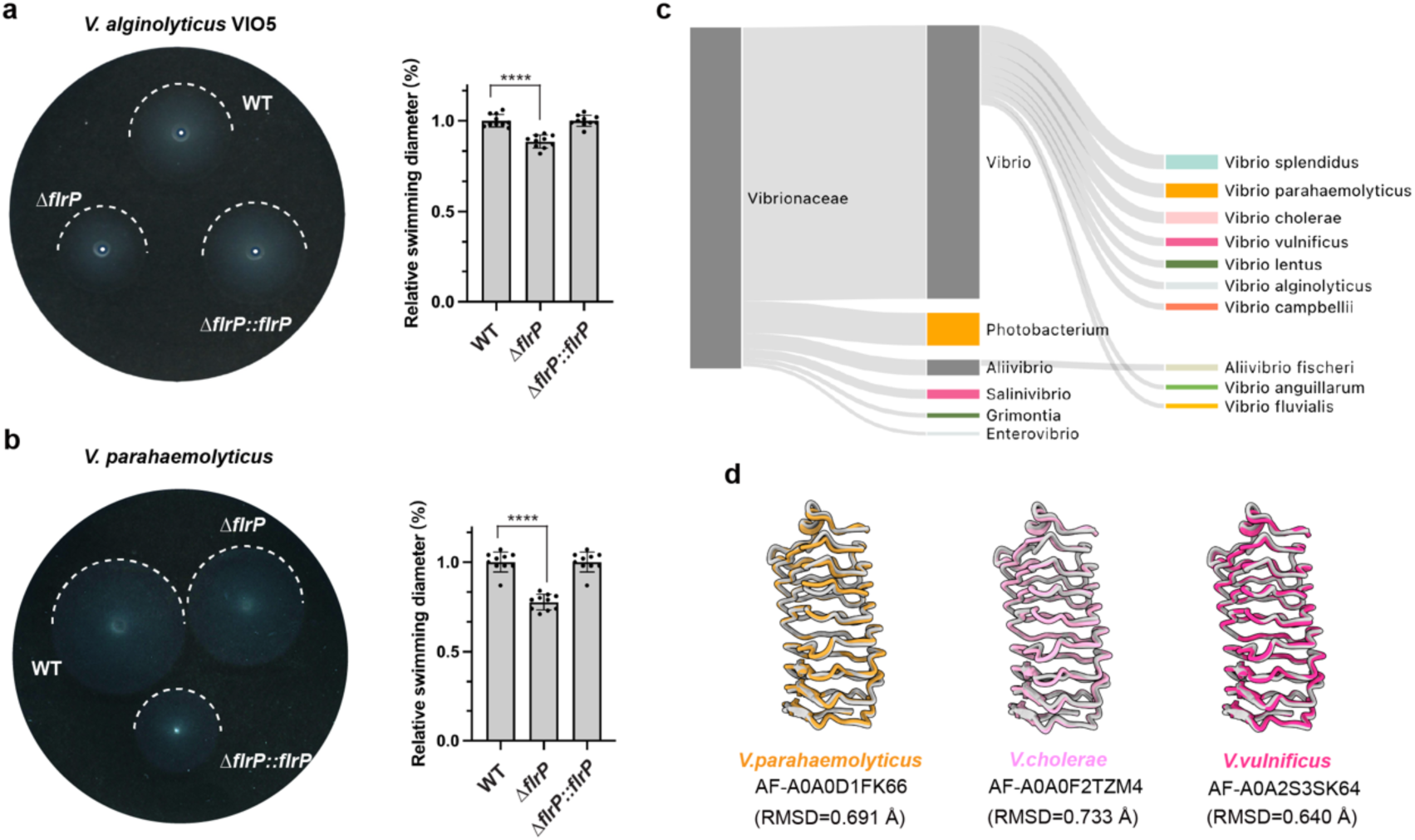
The effect of FlrP in the motility of *V. alginolyticus* and *V. parahaemolyticus*. (a) The effect of gene deletion of *FlrP* on the motility of *V. alginolyticus*. The swimming motility assay of the *V. alginolyticus* VIO5 strain was analyzed using VPG soft-agar plates containing 0.25% agar, quantified after a 6 h incubation at 30℃. Statistical analyses of the effect of gene deletion of *FlrP* on the motility of *V. alginolyticus* are listed at right. (b) The effect of gene deletion of *FlrP* on the motility of *V. parahaemolyticus*. Statistical analyses of the effect are listed at right. The swimming diameters of each sample were measured using ImageJ and normalized to the wild type (WT). Bar graphs represent the mean of at least ten biologically independent replicates with individual data points shown. Statistical significances were determined by a two-tailed Student’s t test (****p < 0.001) (a-b). (c) AFDB-based structural clustering(*49*) of FlrP homologues in *Vibrionaceae*. (d) Structural comparison of the FlrP homologues from *V. parahaemolyticus*, *V. cholerae* and *V. vulnificus*.

**Figure S10:**
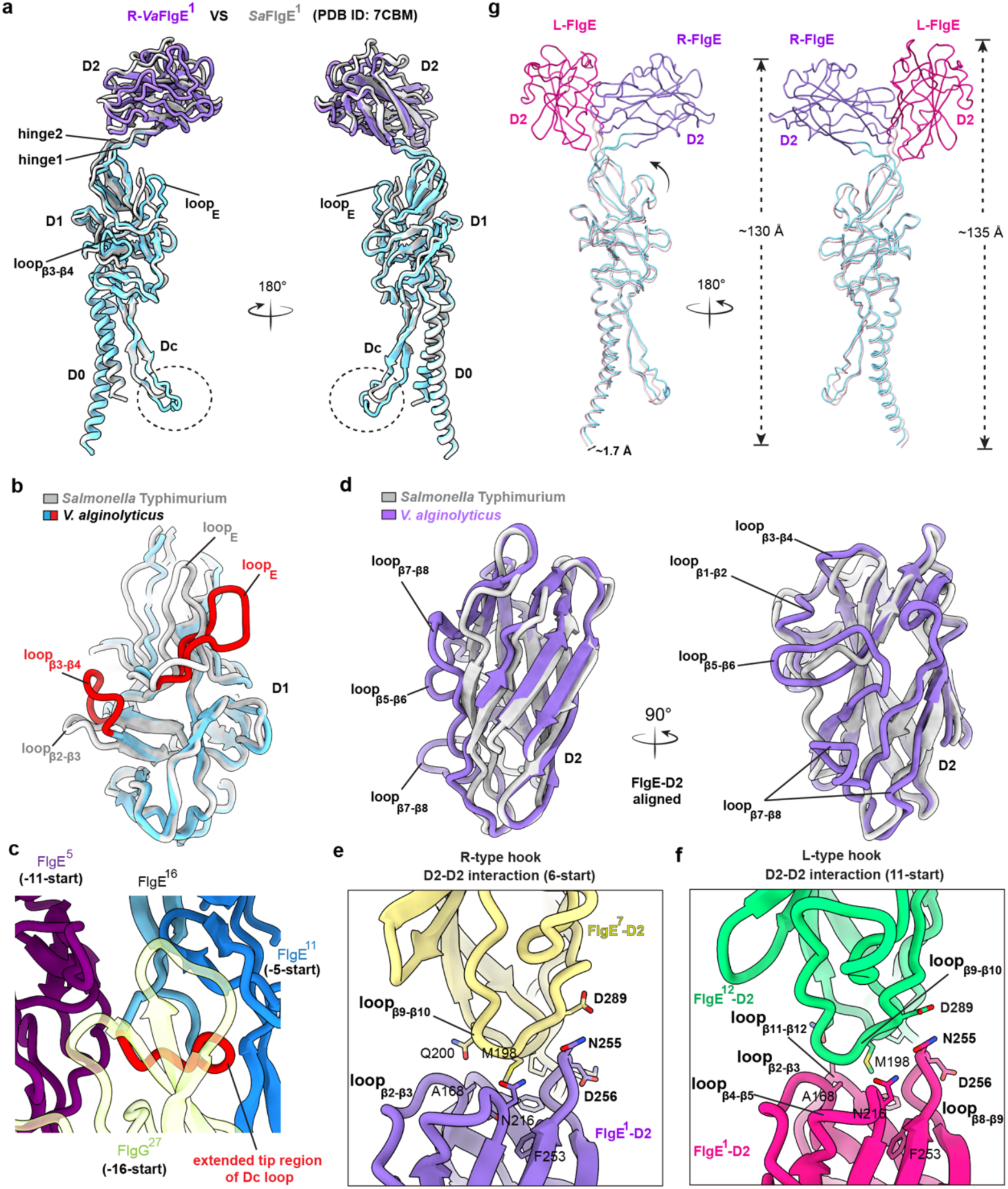
Structural comparison of the FlgE subunits in the R- and L-type hooks. (a) Structural comparison of FlgE of *V. alginolyticus* in the R-type hook (R-*Va*FlgE) of the polar flagellar motor with the *Salmonella* FlgE (*Sa*FlgE) (*14*) of the peritrichous flagellar motor. The tip region is highlighted with dashed circle lines. (b) Structural comparison of the D1 domains of FlgE of *V. alginolyticus* and *Salmonella* Typhimurium. The loop_β3-β4_ and loop_E_ of the D1 domain of the polar FlgE are highlighted in red. (c) The interactions of the Dc loop of the FlgE subunit with the adjacent FlgE and FlgG subunits at the interface of the hook and the rod. (d) Structural comparison of the D2 domains of FlgE of *V. alginolyticus* and *Salmonella* Typhimurium. The distinctive loops in the D2 domains of FlgE of *V. alginolyticus* are labeled as indicated. (e) Detailed interactions of the D2 domains of the R-FlgE subunits along the 6-start direction in the R-type hook. (f) Detailed interactions of the D2 domains of the L-FlgE subunits along the 11-start direction in the L-type hook. (g) Conformational changes of the FlgE subunits in the L- and R-type hooks. The FlgE subunits in the L- and R-type hooks are labeled as L-FlgE and R-FlgE, respectively. The structures of FlgE are shown as ribbons.

**Figure S11:**
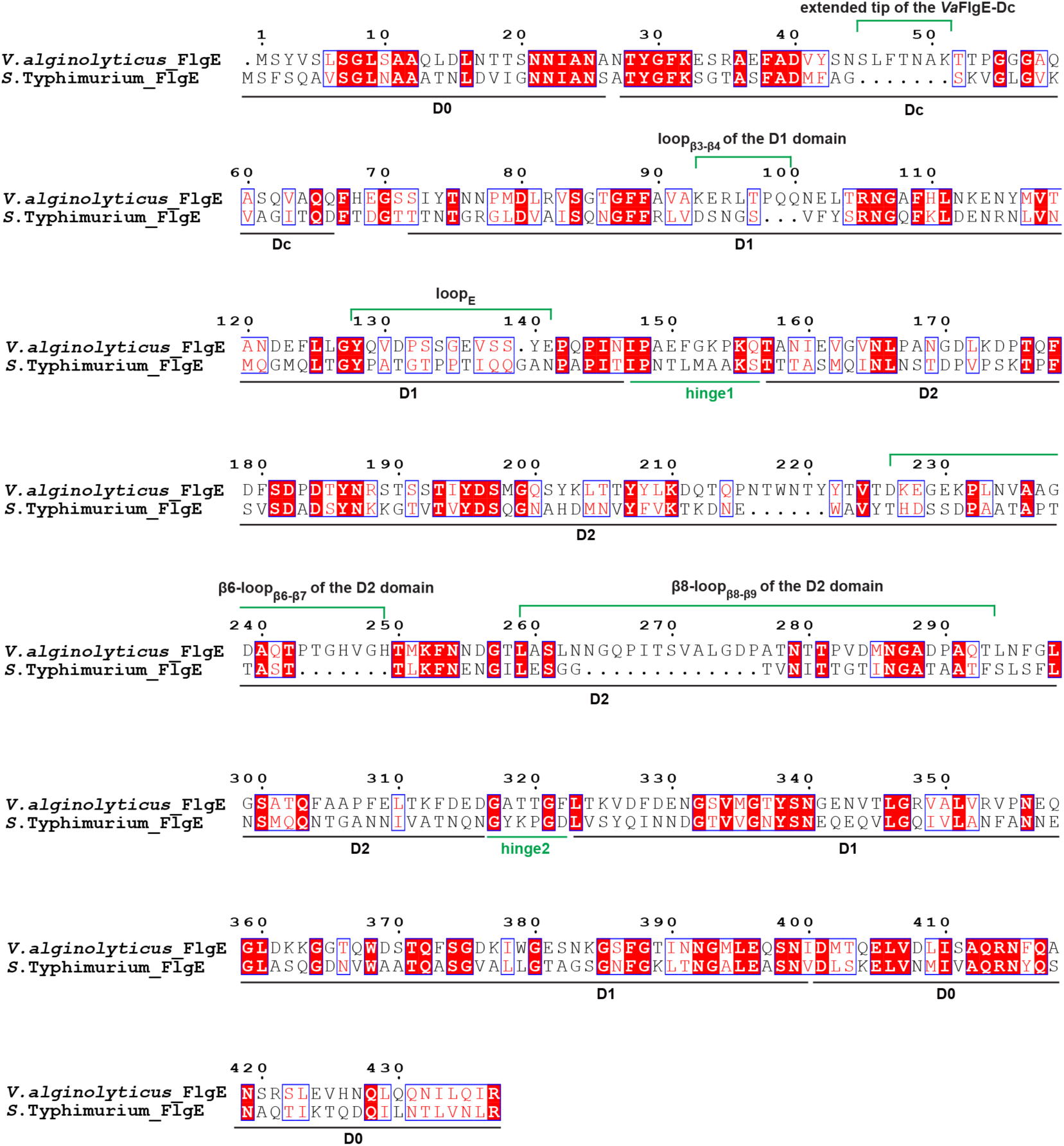
Sequences alignment of the FlgE proteins of *V. alginolyticus* and *S.* Typhimurium. The unique regions in the polar FlgE, including hinge 1 and 2, loop_β3-β4_ and loop_E_ are highlighted as indicated.

**Table S1:** Cryo-EM data collection, refinement and validation statistics.

|  | **LPHT ring**<br>(EMDB-39776)<br>(PDB 8Z5N) | **Proximal rod-**<br>**export apparatus**<br>(EMDB-39780)<br>(PDB 8Z5S) | **Rod-export**<br>**apparatus with**<br>**Partial hook**<br>(EMDB-39782)<br>(PDB 8Z5U) |
| --- | --- | --- | --- |
| **Data collection and processing** |  |  |  |
| Magnification | 105,000 | 105,000 | 105,000 |
| Voltage (kV) | 300 | 300 | 300 |
| Electron exposure (e-/Å<sup>2</sup>) | 50 | 50 | 50 |
| Defocus range (µm) | 1.2 to 1.8 | 1.2 to 1.8 | 1.2 to 1.8 |
| Pixel size (Å) | 1.2 | 1.2 | 1.2 |
| Symmetry imposed | C13 | C1 | C1 |
| Initial particle images (no.) | 329,955 | 329,955 | 329,955 |
| Final particle images (no.) | 45,233 | 58,418 | 58,418 |
| Map resolution (Å) | 2.95 | 3.03 | 3.04 |
| FSC threshold | 0.143 | 0.143 | 0.143 |
| **Refinement** |  |  |  |
| Initial model used (PDB code) | ModelAngelo | ModelAngelo | ModelAngelo |
| Model resolution (Å) | 3.1 | 3.3 | 3.2 |
| FSC threshold | 0.5 | 0.5 | 0.5 |
| Map sharpening *B* factor (Å<sup>2</sup>) | -96.7 | -58.1 | -58.7 |
| Model composition |  |  |  |
| Non-hydrogen atoms | 269,711 | 38,336 | 118,072 |
| Protein residues | 34,671 | 5,016 | 15,597 |
| Ligands | N/A | N/A | N/A |
| *B* factors (Å<sup>2</sup>) |  |  |  |
| Protein | 50.43 | 52.23 | 52.32 |
| Ligand | N/A | N/A | N/A |
| R.m.s. deviations |  |  |  |
| Bond lengths (Å) | 0.004 | 0.003 | 0.003 |
| Bond angles (°) | 0.528 | 0.521 | 0.544 |
| Validation |  |  |  |
| MolProbity score | 1.83 | 1.83 | 1.93 |
| Clashscore | 8.79 | 7.17 | 7.86 |
| Ramachandran plot |  |  |  |
| Favored (%) | 97.90 | 97.94 | 97.56 |
| Allowed (%) | 1.99 | 2.00 | 2.32 |
| Disallowed (%) | 0.11 | 0.06 | 0.12 |

|  | **β-collar-RBM3**<br>**subrings of the**<br>**MS ring (C34)**<br>**(EMDB-39783)**<br>**(PDB 8Z5V)** | **MS ring with the**<br>**proximal rod and**<br>**the export**<br>**apparatus (C1)**<br>**(EMDB-39787)**<br>**(PDB 8Z5X)** | **L-hook**<br>**(EMDB-39788)**<br>**(PDB 8Z5Y)** |
| --- | --- | --- | --- |
| **Data collection and processing** |  |  |  |
| Magnification | 105,000 | 105,000 | 105,000 |
| Voltage (kV) | 300 | 300 | 300 |
| Electron exposure (e-/Å<sup>2</sup>) | 50 | 50 | 50 |
| Defocus range (μm) | 1.2 to 1.8 | 1.2 to 1.8 | 1.2 to 1.8 |
| Pixel size (Å) | 1.2 | 1.2 | 1.2 |
| Symmetry imposed | C34 | C1 | Helical |
| Initial particle images (no.) | 329,955 | 329,955 | 329,955 |
| Final particle images (no.) | 58,418 | 58,418 | 32,950 |
| Map resolution (Å) | 2.95 | 3.64 | 3.22 |
| FSC threshold | 0.143 | 0.143 | 0.143 |
| **Refinement** |  |  |  |
| Initial model used (PDB code) | ModelAngelo | ModelAngelo<br>AlphaFold2 | AlphaFold2 |
| Model resolution (Å) | 3.2 | 6.2 | 3.5 |
| FSC threshold | 0.5 | 0.5 | 0.5 |
| Map sharpening *B* factor (Å<sup>2</sup>) | -109.9 | -52.1 | -77.7 |
| Model composition |  |  |  |
| Non-hydrogen atoms | 39,814 | 88,328 | 132,760 |
| Protein residues | 5,032 | 11,398 | 17,400 |
| Ligands | N/A | N/A | N/A |
| *B* factors (Å<sup>2</sup>) |  |  |  |
| Protein | 44.26 | 58.09 | 65.49 |
| Ligand | N/A | N/A | N/A |
| R.m.s. deviations |  |  |  |
| Bond lengths (Å) | 0.003 | 0.004 | 0.005 |
| Bond angles (°) | 0.479 | 0.563 | 0.613 |
| Validation |  |  |  |
| MolProbity score | 1.88 | 2.31 | 2.05 |
| Clashscore | 5.21 | 9.75 | 6.22 |
| Ramachandran plot |  |  |  |
| Favored (%) | 97.18 | 96.05 | 96.65 |
| Allowed (%) | 2.82 | 3.71 | 3.22 |
| Disallowed (%) | 0.00 | 0.24 | 0.13 |

|  | **R-hook**<br>(EMDB-39789)<br>(PDB 8Z5Z) | **LRR-rod-FliF**<br>**Peptide loops**<br>(EMDB-39786)<br>(PDB 8Z5W) | **The polar**<br>**flagellar**<br>**motor-hook**<br>**complex**<br>(EMDB-39790)<br>(PDB 8Z60) |
| --- | --- | --- | --- |
| **Data collection and processing** |  |  |  |
| Magnification | 105,000 | 105,000 | 105,000 |
| Voltage (kV) | 300 | 300 | 300 |
| Electron exposure (e-/Å<sup>2</sup>) | 50 | 50 | 50 |
| Defocus range (μm) | 1.2 to 1.8 | 1.2 to 1.8 | 1.2 to 1.8 |
| Pixel size (Å) | 1.2 | 1.2 | 1.2 |
| Symmetry imposed | Helical | C1 | C100 |
| Initial particle images (no.) | 329,955 | 329,955 | 329,955 |
| Final particle images (no.) | 19,402 | 19,450 | 61,395 |
| Map resolution (Å) | 3.42 | 3.46 | 3.22 |
| FSC threshold | 0.143 | 0.143 | 0.143 |
| **Refinement** |  |  |  |
| Initial model used (PDB code) | AlphaFold2 | ModelAngelo |  |
| Model resolution (Å) | 3.9 | 3.7 |  |
| FSC threshold | 0.5 | 0.5 |  |
| Map sharpening *B* factor (Å<sup>2</sup>) | -61.2 | -44.8 |  |
| Model composition |  |  |  |
| Non-hydrogen atoms | 132,760 | 36,128 |  |
| Protein residues | 17,400 | 4,810 |  |
| Ligands | N/A | N/A |  |
| *B* factors (Å<sup>2</sup>) |  |  |  |
| Protein | 65.49 | 54.19 |  |
| Ligand | N/A | N/A |  |
| R.m.s. deviations |  |  |  |
| Bond lengths (Å) | 0.002 | 0.003 |  |
| Bond angles (°) | 0.416 | 0.525 |  |
| Validation |  |  |  |
| MolProbity score | 1.75 | 1.92 |  |
| Clashscore | 5.84 | 7.64 |  |
| Ramachandran plot |  |  |  |
| Favored (%) | 97.74 | 97.24 |  |
| Allowed (%) | 2.06 | 2.63 |  |
| Disallowed (%) | 0.00 | 0.13 |  |

